# TBK1 eliminates aggregation-prone monomeric TDP-43 through the IFNβ-immunoproteasome pathway

**DOI:** 10.1101/2025.04.29.650889

**Authors:** Shohei Sakai, Kotaro Oiwa, Yohei Iguchi, Seiji Watanabe, Okiru Komine, Mai Horiuchi, Masahisa Katsuno, Koji Yamanaka

## Abstract

Loss-of-function mutations in TANK-binding kinase 1 (TBK1) are genetically linked to amyotrophic lateral sclerosis (ALS) and frontotemporal dementia (FTD), and induce cytoplasmic aggregation of TAR DNA-binding protein 43 (TDP-43), known as TDP-43 pathology. Although TBK1 deficiency is thought to contribute to TDP-43 pathology primarily through impaired autophagy, the full spectrum of its pathological impact remains unclear. Given the multifunctional nature of TBK1, alternative pathways beyond autophagy are possibly involved in TDP-43 pathology. Here, we found that TBK1 alleviates TDP-43 pathology in neuronal cells via induction of interferon-beta (IFNβ), and that the IFNβ receptor is downregulated in spinal motor neurons from ALS patients with TDP-43 pathology. We further demonstrated that IFNβ induces the immunoproteasome by upregulating its subunits, thereby promoting the degradation of aggregation-prone monomeric TDP-43. Furthermore, heterozygous deletion of *Tbk1* in *SOD1*^G93A^ ALS model mice resulted in reduced immunoproteasome induction and increased polyubiquitinated protein accumulation in the spinal cord. These findings suggest that impairment of the TBK1-IFNβ-immunoproteasome axis may contribute to the development of TDP-43 pathology in ALS and FTD.

## Introduction

Cytoplasmic mislocalization and abnormal accumulation of TAR DNA-binding protein 43 (TDP-43) in the brain and spinal cord, known as TDP-43 pathology, are pathological hallmarks in several incurable neurodegenerative diseases such as amyotrophic lateral sclerosis (ALS) and frontotemporal dementia (FTD) (Arai *et al*, 2006; Neumann *et al*, 2006). TDP-43 is an RNA-binding protein that regulates multiple aspects of RNA metabolism such as RNA transcription, splicing, transport, and translation (Lagier-Tourenne *et al*, 2010). As TDP-43 regulates numerous transcripts essential for neuronal health and function (Polymenidou *et al*, 2011; Klim *et al*, 2019; Melamed *et al*, 2019; Brown *et al*, 2022; Ma *et al*, 2022), the cytoplasmic aggregation of TDP-43, along with the loss of nuclear TDP-43, is considered a primary driver of neurodegeneration. Therefore, preventing TDP-43 pathology represents a rational therapeutic strategy for ALS and FTD. However, the precise molecular mechanisms underlying TDP-43 pathology remain incompletely understood.

*TBK1* is a potentially important gene in the context of TDP-43 pathology because mutations in *TBK1* cause familial ALS/FTD with TDP-43 pathology (Cirulli *et al*, 2015; Freischmidt *et al*, 2015) and its gene product, TANK-binding kinase 1 (TBK1), is involved in the autophagy (Oakes *et al*, 2017) and the endosome pathway (Hao *et al*, 2021; Shao *et al*, 2022), both of which contribute to the degradation of large protein aggregate depending on lysosomal activity. Additionally, we recently reported that the level of active TBK1, phosphorylated TBK1, is reduced in sporadic ALS patients’ brains and spinal cords (Watanabe *et al*, 2023), suggesting that TBK1 has important roles not only in familial but also in sporadic ALS/FTD cases.

TBK1 is a multifunctional kinase involved in diverse cellular events such as autophagy, programmed cell death, and inflammation (Oakes *et al*, 2017; Xu *et al*, 2018; Ahmad *et al*, 2016), and loss of function in *TBK1* due to mutations is considered a responsible mechanism for pathogenesis of familial ALS/FTD (Freischmidt *et al*, 2015).

In the autophagy pathway, TBK1 phosphorylates autophagy receptors Optineurin (Formstone *et al*, 2011) and p62 (Pilli *et al*, 2012) and promotes autophagic clearance of damaged organelles and protein aggregates. Because *OPTN* and *SQSTM1* have been commonly identified as ALS/FTD causative genes (Maruyama *et al*, 2010; Fecto *et al*, 2011; Le Ber *et al*, 2013; Pottier *et al*, 2018), there is growing consensus that dysregulation of the autophagy-lysosome pathway is one of the main causes of ALS/FTD pathogenesis and TDP-43 pathology. Nonetheless, previous studies using single genetic manipulations of *Tbk1*, such as systemic heterozygous knockout, motor neuron-specific conditional knockout, and knock-in of ALS/FTD-linked mutations, were insufficient to recapitulate the disease phenotypes and TDP-43 pathology in mice (Brenner *et al*, 2019; Gerbino *et al*, 2020; Sieverding *et al*, 2021).

In innate immune signaling, TBK1 regulates inflammation by promoting cytokine secretion. TBK1 phosphorylates stimulator of interferon genes (STING), nuclear factor-kappa B (NF-κB), and interferon regulatory factor 3 (IRF3) to induce the expression of pro-inflammatory cytokines such as tumor necrosis factor (TNF) and type 1 interferons (IFNs) such as IFNβ. Recent studies have suggested that blocking the TBK1-associated innate immune signaling pathway by STING suppression ameliorates disease progression of ALS model mice and induced pluripotent stem cell (iPSC)-derived motor neurons (Yu *et al*, 2020; Tan *et al*, 2022). However, these findings appear inconsistent with the concept that the loss of function mutations in *TBK1* causes familial ALS/FTD. Besides, while inflammatory cytokines and IFNs are well-known as modulators of glial activity and functions, their direct effect on ALS/FTD-susceptible neurons and TDP-43 pathology have not been fully investigated.

Here, we propose a novel mechanism of abnormal TDP-43 clearance by TBK1. In this study, we found that TBK1-induced IFNβ promotes the degradation of aggregation-prone TDP-43 via the immunoproteasome pathway in neuronal cells. Importantly, the receptor for IFNβ was downregulated in spinal motor neurons of ALS patients. In spinal cords and cerebral cortices of mice, immunoproteasome induction was observed in aged wild-type mice and *SOD1*^G93A^ ALS model mice. Heterozygous deletion of *Tbk1* in the ALS mice resulted in reduced immunoproteasome induction and enhanced accumulation of polyubiquitinated proteins at onset stage. These findings suggest, for the first time, that impairment of the TBK1-IFNβ-immunoproteasome axis may contribute to the pathogenesis of ALS/FTD associated with TDP-43 pathology.

## Results

### TBK1 reduces aggregation-prone monomeric TDP-43 via humoral factors

We recently reported that the monomerization of TDP-43 is a key determinant of TDP-43 pathology in sporadic ALS cases, and that expression of monomeric TDP-43 mutants in neuronal cells recapitulates key features of TDP-43 pathology (Oiwa *et al*, 2023). To examine the effect of TBK1 function on TDP-43 pathology, we utilized a cell-based TDP-43 pathology model using a monomeric TDP-43 mutant named 6M (TDP-43^6M^) (Afroz *et al*, 2017) and induced TBK1 overexpression in the system. Neuro2a cells co-transfected with mock or human Myc-TBK1 and human TDP-43^WT^ or TDP-43^6M^-3xFLAG constructs were fractionated into 1% Triton X-100-soluble and -insoluble fractionations, followed by immunoblotting analysis (Fig. 1A). TBK1 overexpression reduced TDP-43 levels in both the soluble and insoluble fractions, with a more pronounced effect on TDP-43^6M^ than on TDP-43^WT^. TBK1 also decreased the levels of phosphorylated TDP-43 (pTDP) induced by the monomeric mutant. We confirmed that TBK1 overexpression did not affect the mRNA levels of the transfected TDP-43 (Fig. S1). In contrast to TBK1 overexpression, reduction of endogenous TBK1 activity via siRNA-mediated knockdown or treatment with a TBK1-specific inhibitor BX795 resulted in an increase in pTDP levels (Fig. S2). The reduction of TDP-43^6M^ by TBK1 overexpression was dependent on its kinase activity, as ALS/FTD-linked kinase dead TBK1 mutants failed to reduce TDP-43^6M^ levels (Fig. 1B). As TBK1 is known to be involved in autophagy, we initially hypothesized that the marked reduction of the monomeric TDP-43 mutant was dependent on the autophagy–lysosome pathway. To test this hypothesis, Neuro2a cells co-transfected with TDP-43^6M^ and TBK1 were treated with bafilomycin A1, a potent inhibitor of the autophagy–lysosome pathway. While TBK1 increased the levels of phosphorylated p62 and LC3-II, supporting the activation of autophagy, TDP-43^6M^ levels were still reduced by TBK1 even under the inhibitor treatment (Fig. 1C). These results suggested that TBK1 has the potential to ameliorate TDP-43 pathology by reducing excess TDP-43, particularly aggregation-prone monomeric TDP-43, through a mechanism that is independent of the autophagy–lysosome pathway. Next, we investigated the possibility that TBK1 suppresses TDP-43 pathology via paracrine signaling, as TBK1 is known to be involved in cytokine secretion. Interestingly, co-culture experiments illustrated in Figure 1D, followed by immunoblotting (Fig. S3A) and immunocytochemistry (Fig. 1E), showed that protein levels of TDP-43^6M^ and pTDP were reduced by neighboring cells overexpressing TBK1. To rule out the effects of physical cell–cell contact in the co-culture system, we conducted an additional experiment using conditioned medium (CM) (Fig. 1F). As expected, CM derived from TBK1-overexpressing cells also reduced levels of TDP-43^6M^ and pTDP (Fig. 1G and Fig. S3B). Collectively, these findings suggested that TBK1 ameliorates TDP-43 pathology via humoral factors such as cytokines.

**Figure 1.**
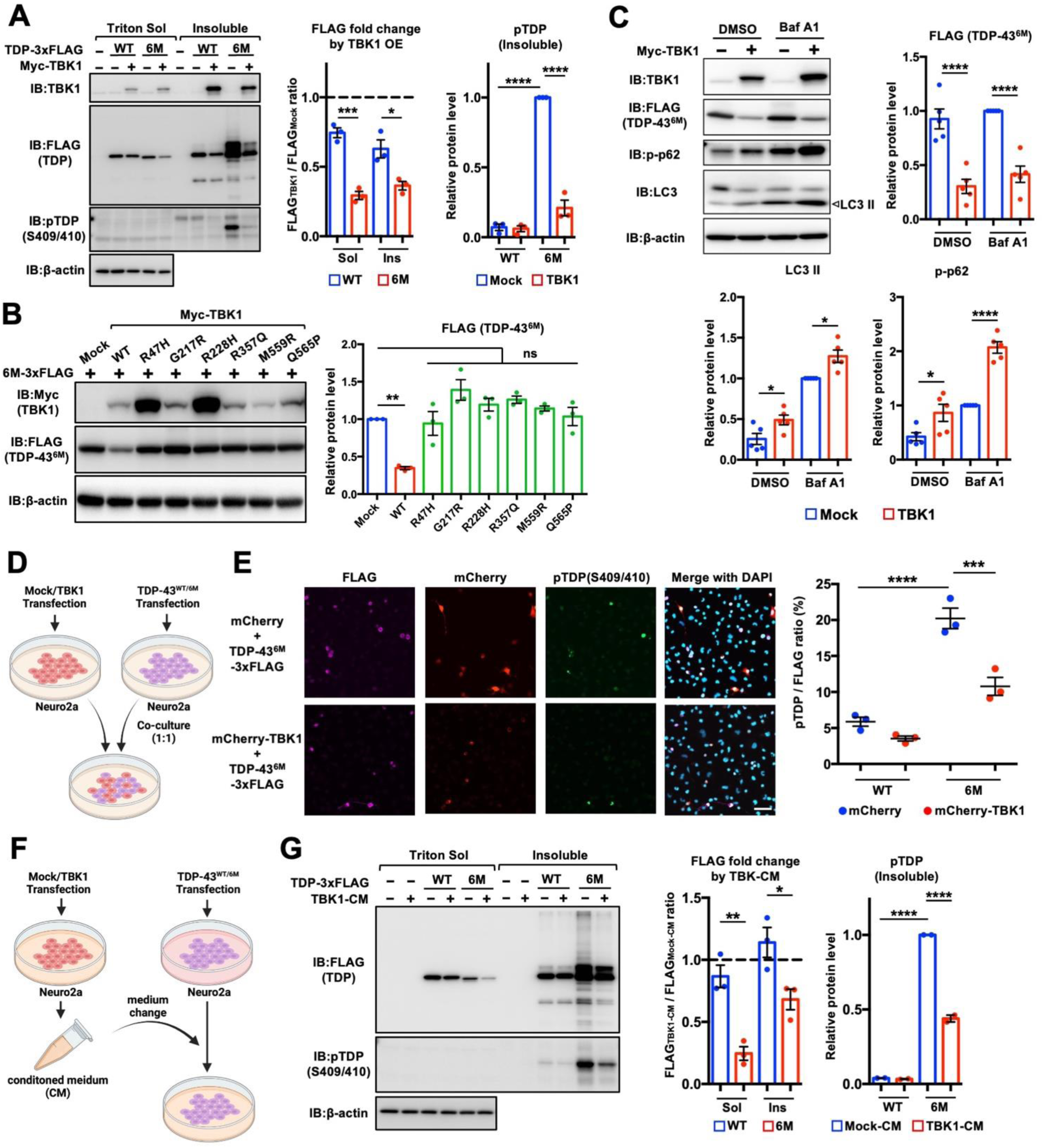
TBK1 overexpression ameliorates TDP-43 pathology in cells via humoral factors. **A.** TBK1 overexpression reduced the levels of transfected human TDP-43, particularly the monomeric TDP-43 mutant, as well as its phosphorylated form (pTDP). Neuro2a cells were transiently co-transfected with human TDP-43^WT^ or TDP-43^6M^-3×FLAG and either mock or human Myc-TBK1 expression vectors, then fractionated into 1% Triton X-100-soluble and -insoluble fractions and subjected to immunoblot analysis. **B.** ALS/FTD-linked kinase-dead TBK1 mutants failed to reduce monomeric TDP-43 levels. Neuro2a cells were transiently co-transfected with TDP-43^6M^-3×FLAG and the indicated mutant expression vectors of Myc-TBK1. **C.** Inhibition of the autophagy–lysosome pathway did not abolish the TBK1-mediated reduction of monomeric TDP-43. Neuro2a cells were transiently co-transfected with TDP-43^6M^-3×FLAG and either mock or Myc-TBK1 expression vectors, treated with either DMSO or 100 nM Bafilomycin A1 (Baf A1) for 20 hours, and analyzed by immunoblotting. **D, E.** TDP-43 pathology was suppressed by neighboring cells overexpressing TBK1. The experimental scheme is illustrated in (D). Representative immunofluorescence images of co-cultured Neuro2a are shown in (E). Only images of the TDP-43^6M^ case are shown. The ratio of pTDP-positive to FLAG-positive cells is quantified in the graph. Scale bar, 50 μm. **F, G.** Conditioned medium (CM) from TBK1-overexpressing cells reduced monomeric TDP-43 and pTDP levels in CM-recipient cells. The experimental scheme is illustrated in (F). CM-recipient cells were fractionated as with A and analyzed by immunoblotting. Data are presented as means ± SEM. ns, not significant; *p < 0.05, **p < 0.01, ***p < 0.001, ****p < 0.0001.

### TBK1-IFN**β** pathway is crucial for clearance of monomeric TDP-43

In innate immunity, TBK1 mediates cytokine secretion through phosphorylation of STING and IRF3 (Shu *et al*, 2014; Liu *et al*, 2015). To test whether the observed humoral factor-mediated reduction of monomeric TDP-43 depends on this innate immune pathway, we pretreated cells with siRNAs targeting *Sting* or *Irf3* prior to TBK1 overexpression and again performed the experiment using CM (Fig. 2A). qPCR analysis of the CM-donor cells confirmed efficient knockdown of both *Sting* and *Irf3*, and that these treatments did not affect the expression level of transfected human TBK1 (Fig. 2B). Immunoblotting of the CM-recipient cells showed that knockdown of either Sting or Irf3 abolished the reduction of TDP-43^6M^ and pTDP levels by TBK1-CM (Fig. 2C). Because we observed strong induction of *Ifnb1* (IFNβ) gene expression upon TBK1 overexpression (Fig. 2D), which was significantly suppressed by knockdown of *Sting* or *Irf3* (Fig. 2E), we next focused on IFNβ. We prepared two distinct siRNAs targeting *Ifnb1* and confirmed that both effectively inhibited TBK1-induced *Ifnb1* expression (Fig. 2F). As expected, immunoblotting of the CM-recipient cells revealed that knockdown of *Ifnb1* in CM-donor cells completely abolished the effect of TBK1-CM on TDP-43^6M^ and pTDP levels, demonstrating that IFNβ is the key cytokine responsible for TBK1-mediated clearance of TDP-43^6M^. We also confirmed that inhibition of the JAK/STAT pathway, which is a canonical downstream signaling pathway of IFNβ, by a pan-JAK inhibitor tofacitinib completely abolished the reduction of TDP-43^6M^ by TBK1-CM. Collectively, these findings demonstrate that the TBK1-STING-IRF3-IFNβ-JAK-STAT axis is crucial for the clearance of monomeric TDP-43. The increase in pTDP observed upon suppression of endogenous TBK1 (Fig. S2) is unlikely to depend on the same pathway, as siTbk1 and BX795 treatment did not affect *Ifnb1* mRNA levels (Fig. S4).

**Figure 2.**
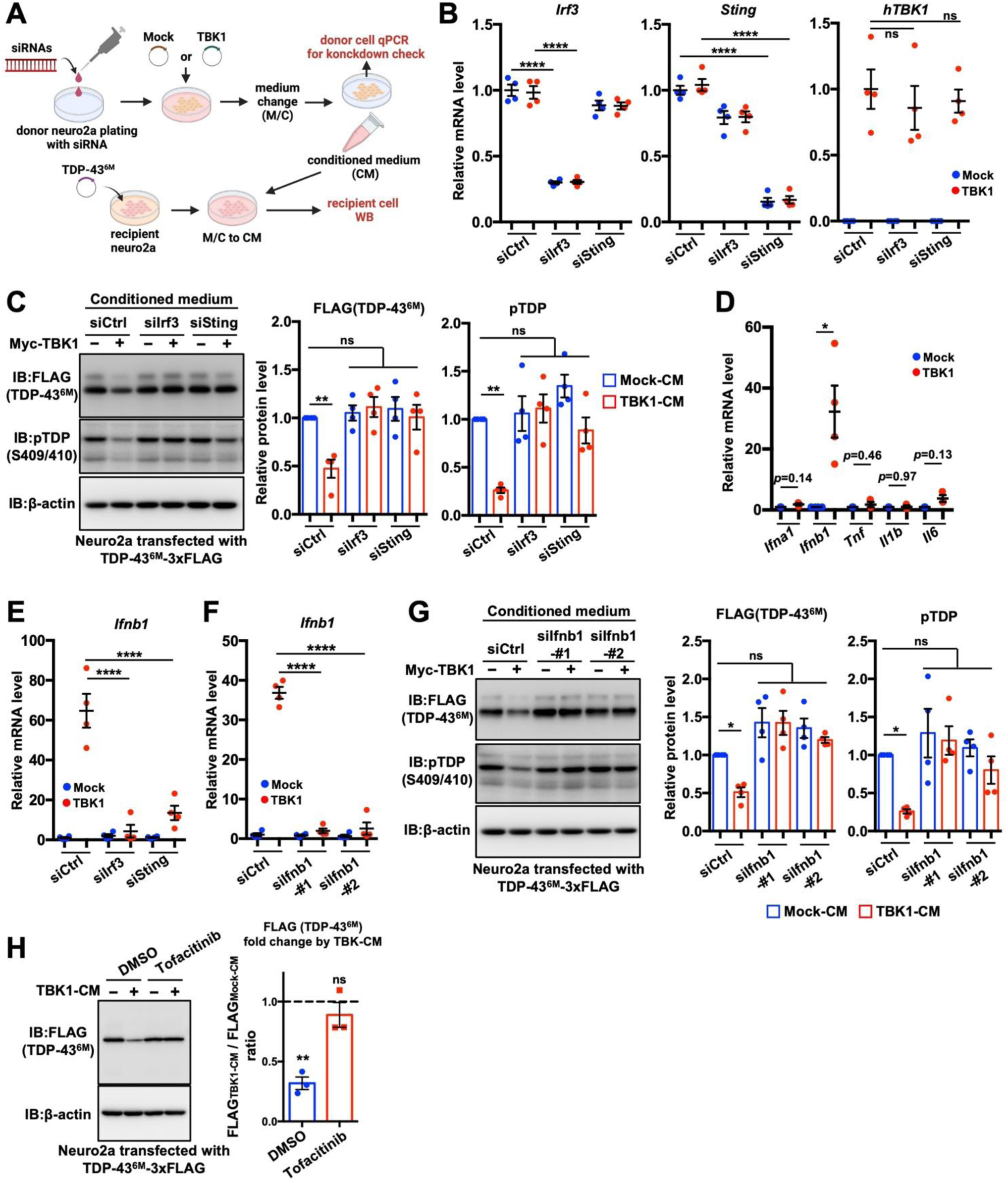
The STING-IRF3-IFNβ pathway ameliorates TDP-43 pathology in cells. **A.** A schematic illustration of experiments using siRNAs and conditioned media (CM) performed in this figure. **B, C.** Knockdown of *Irf3* or *Sting* abolished the reduction of the monomeric TDP-43 by TBK1-CM. Efficient reduction of *Irf3* and *Sting* mRNA expression, without affecting *hTBK1*, in CM-donor cells was confirmed by qPCR (B). Cells treated with the indicated CM conditions were analyzed by immunoblotting (C). **D.** TBK1 overexpression in Neuro2a cells markedly induced IFNβ (*Ifnb1*). Expression levels of the indicated genes were analyzed by qPCR. **E.** qPCR revealed that knockdown of either *Sting* or *Irf3* markedly suppressed *Ifnb1* induction by TBK1 in Neuro2a cells. **F, G.** Knockdown of *Ifnb1* abolished the reduction of the monomeric TDP-43 by TBK1-CM. Significant reduction of *Ifnb1* expression in CM-donor cells was confirmed by qPCR (F). Cells treated with the indicated CM conditions were analyzed by immunoblotting (G). (H) Pan-JAK inhibition abolished the reduction of the monomeric TDP-43 by TBK1-CM. Neuro2a cells transiently transfected with TDP-43^6M^-3xFLAG were treated with either Mock-or TBK1-CM in the presence of either DMSO or 5 μM Tofacitinib. Data are expressed as means ± SEM. ns, not significant. *p < 0.05, **p < 0.01, and ****p < 0.0001.

### IFN**β** treatment reduces monomeric TDP-43 in neuronal cells

Next, we tested whether IFNβ treatment alone is sufficient to reduce TDP-43 levels. Neuro2a cells were transfected with TDP-43^WT^ or its mutants, including another monomeric TDP-43 mutant L27/28A (Mompeán *et al*, 2017) and nuclear localization signal-deficient mutant (*Δ*NLS) (Winton *et al*, 2008). The cells were then treated with recombinant mouse IFNβ and subjected to detergent solubility fractionation followed by immunoblot analysis. IFNβ treatment significantly reduced both monomeric TDP-43 mutants in the 1% Triton X-100-soluble and -insoluble fractions, as well as pTDP in the insoluble fraction (Fig. 3A). Immunocytochemistry further confirmed that recombinant IFNβ treatment decreased the number of pTDP-positive cells (Fig. 3B). We also generated Neuro2a and NSC-34 cell lines stably expressing doxycycline-inducible TDP-43^WT^ or TDP-43^6M^-3xFLAG and confirmed that IFNβ treatment selectively reduced TDP-43^6M^ levels (Fig. 3C and S5). Similar results were obtained in the human neuronal cell line SH-SY5Y (Fig. 3D), but not in HEK293 cells (Fig. 3E). Additionally, we found that ALS-linked SOD1 mutants were also reduced by IFNβ (Fig. S6). Collectively, these findings suggest that IFNβ broadly reduces aggregation-prone proteins, and that this regulatory mechanism may be specific to neuronal cells.

**Figure 3.**
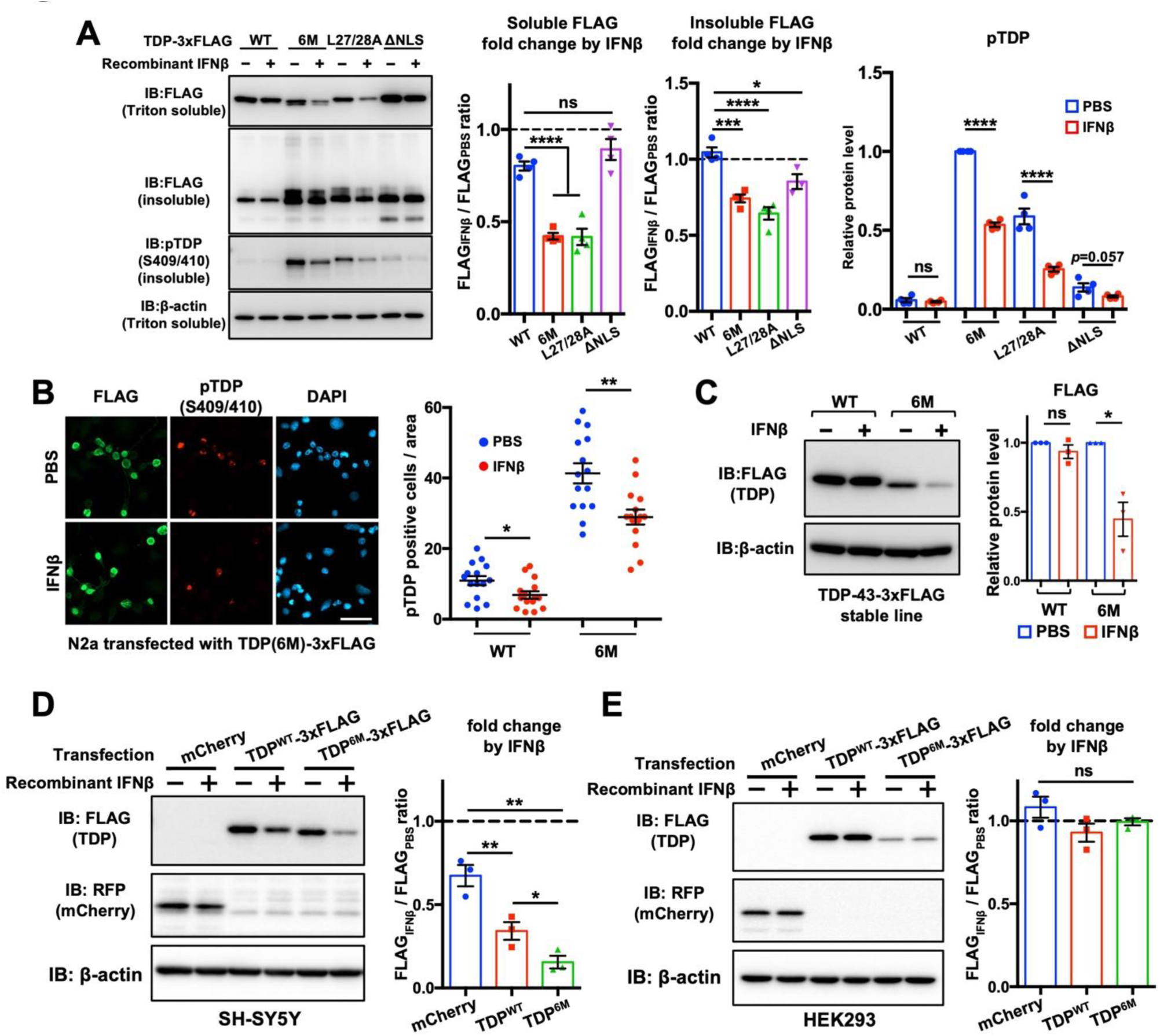
IFNβ treatment reduces monomeric TDP-43 in neuronal cells. **A, B.** Recombinant IFNβ treatment reduced monomeric TDP-43 mutants and suppressed pTDP accumulation. Neuro2a cells transiently transfected with expression vectors of the indicated TDP-43 mutants were treated with either PBS or 10 ng/ml mouse IFNβ for 42h, fractionated into 1% TritonX-100-soluble and -insoluble fractions, and analyzed by immunoblotting (A). Reduction of pTDP-positive cells by recombinant IFNβ was confirmed by immunocytochemistry (B). Only images of the TDP-43^6M^ case are shown. The number of pTDP-positive cells in the counted area is presented in the graph. Scale bar, 50 μm. **C.** Recombinant IFNβ selectively reduced the monomeric TDP-43 mutant in stable lines. Neuro2a cells stably expressing Dox-inducible TDP-43^WT^ or TDP-43^6M^-3xFLAG were treated with 1.5 μg/ml Dox and either PBS or 10 ng/ml mouse IFNβ for 48h and analyzed by immunoblotting. **D, E.** Recombinant IFNβ treatment reduced TDP-43 in SH-SY5Y but not in HEK293. SH-SY5Y cells (D) and HEK293 cells (E) were transiently transfected with the indicated expression vectors, treated with either PBS or 10ng/mL human IFNβ for 48h, and analyzed by immunoblotting. Data are expressed as means ± SEM. ns, not significant. *p < 0.05, **p < 0.01***, p < 0.001, and ****p < 0.0001.

### IFN**β** receptor is downregulated in spinal motor neurons of ALS patients

IFNβ binds to interferon α/β receptor (IFNAR), a heterodimer composed of IFNAR1 and IFNAR2, to activate downstream signaling (Ivashkiv & Donlin, 2014). Although IFNAR is reported to be ubiquitously expressed based on studies using systemic IFNAR1-null mice (Müller *et al*, 1994) and the integrated single-cell/nucleus RNA-sequencing data from multiple tissues in the Human Protein Atlas (http://www.proteinatlas.org/celltype) (Karlsson *et al*, 2021) (Fig. S7), its detailed protein-level expression pattern in the spinal cord and brain has not been fully characterized. In this study, we investigated whether ALS/FTD-susceptible neurons express the receptor for IFNβ by performing immunostaining for IFNAR1, one of the two subunits of the receptor. Immunofluorescence analysis of IFNAR1 in the lumbar spinal cord and cerebral cortex of wild-type mice revealed that it is primarily expressed in NeuN-positive neurons, including choline acetyltransferase (ChAT)-positive spinal motor neurons and COUP-TF-interacting protein 2 (CTIP2)-positive layer V cortical neurons (Fig. 4A, B). These findings suggest that type I IFN signaling plays important roles in neurons, possibly including IFNβ-mediated clearance of aggregation-prone proteins. Next, we investigated whether IFNAR1 protein expression is altered in motor neurons of sporadic ALS patients with TDP-43 pathology. In the controls, strong IFNAR1 signals were observed in motor neurons, consistent with the results obtained in mice (Fig. 4C). However, the fluorescence intensity in motor neurons from ALS patients (Fig. 4D) was significantly lower than that in the controls (Fig. 4E and S8). Furthermore, we compared IFNAR1 fluorescence intensity between TDP-43 pathology-negative and -positive motor neurons within ALS patients and found that IFNAR1 expression was lower in pathology-positive neurons (Fig. 4F). These findings suggest that dysregulated IFNβ signaling in motor neurons may be associated with TDP-43 pathology in ALS.

**Figure 4.**
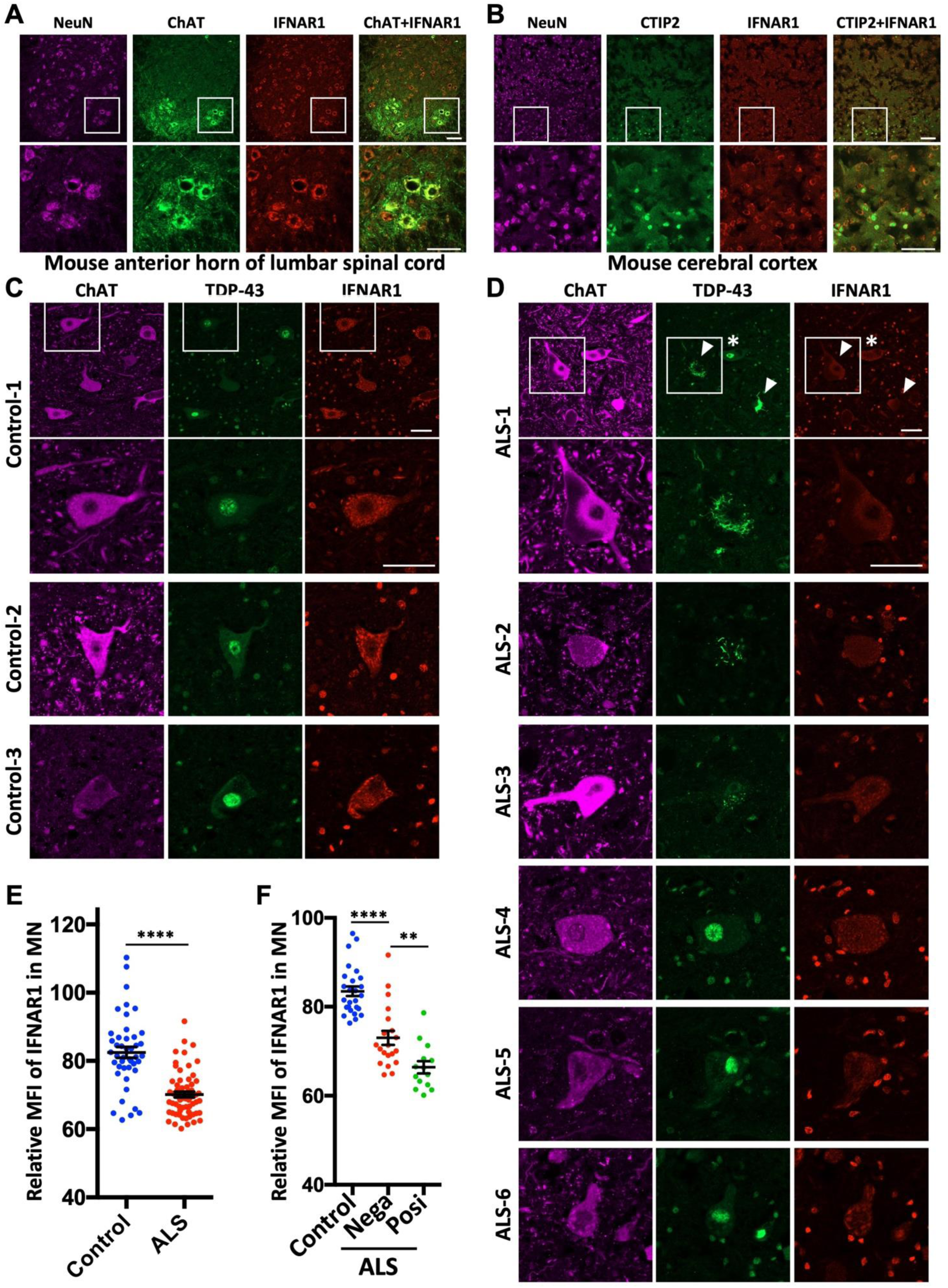
IFNAR1, a subunit of the receptor for IFNβ, is expressed primarily in neurons, but the expression level is reduced in ALS patients’ spinal motor neurons. **A, B.** IFNAR1 is expressed primarily in neurons. Representative images of IFNAR1 immunostaining with a pan-neuron marker NeuN and either a spinal motor neuron marker ChAT or a layer V cortical neuron marker CTIP2 in lumbar spinal cords (A) or cerebral cortices (B) of WT mice. Scale bar, 100 μm. **C-F.** IFNAR1 is downregulated in spinal motor neurons of ALS patients. Representative images of IFNAR1 immunostaining with ChAT and TDP-43 in spinal cords of controls (C) and ALS patients (D). The asterisk and arrowheads indicate TDP-43 pathology-negative and -positive motor neurons, respectively. Scale bar, 50 μm. Relative mean fluorescent intensities (MFI) of IFNAR1 in a single motor neuron are shown in E and F. In F, results of motor neurons from 2 controls (Control-1 and -2) and 3 ALS patients (ALS-1, -2, and -3) are shown, in which motor neurons from ALS patients are subdivided into two groups based on the presence or absence of TDP-43 pathology. MN, motor neuron. Nega, TDP-43 pathology-negative. Posi, TDP-43 pathology-Positive. Data are expressed as means ± SEM. **p < 0.01 and ****p < 0.0001.

### IFN**β** promotes degradation of monomeric TDP-43 through the immunoproteasome pathway

Next, we explored the downstream mechanisms of IFNβ. In our previous results, IFNβ induced the reduction of not only the aggregated form but also the mild detergent-soluble form of monomeric TDP-43 mutants (Fig. 1G and 3A), which are typically degraded via the ubiquitin– proteasome pathway. Therefore, we focused on this degradation pathway and found that treatment with canonical proteasome inhibitors, MG132 or lactacystin, completely abolished the IFNβ-induced reduction of TDP-43^6M^ (Fig. 5A, B). Given this proteasome dependency, we next examined whether the immunoproteasome, a subtype of the proteasome, might contribute to the clearance of monomeric TDP-43. The immunoproteasome is a specialized form of the proteasome complex normally induced by proinflammatory cytokines, such as TNF and IFN*γ*, particularly in the context of viral infection. Its overall structure is similar to that of the canonical proteasome, but it incorporates two immunoproteasome-specific regulatory subunits, PSME1 and PSME2 (also known as PA28α and PA28β, respectively), and three catalytic β-subunits, PSMB8, PSMB9, and PSMB10, thereby facilitating the generation of peptides for MHC class I antigen presentation (McCarthy & Weinberg, 2015) (Fig. 5C). Although IFNβ is not considered a canonical inducer of the immunoproteasome, previous studies have shown that type I IFNs can also induce immunoproteasome subunits in hepatocytes (Shin *et al*, 2006), epithelial cells, and fibroblasts (Wang *et al*, 2023). We therefore tested whether IFNβ upregulates immunoproteasome subunits in neurons and found that IFNβ robustly induced PSMB8 and PSMB9 in three neuronal cell lines (SH-SY5Y, Neuro2a, and NSC-34 cells), but not in HEK293 or HeLa cells (Fig. 5D). qPCR analysis of Neuro2a cells also revealed that both IFNβ and TBK1-CM induced the expression of catalytic β-subunits and regulatory subunits of the immunoproteasome (Fig. 5E and 5F). Next, we assessed immunoproteasome activity using a PSMB8-specific fluorogenic substrate, Ac-Ala-Asn-Trp-7-amino-4-methylcoumarin (Ac-ANW-AMC). As expected, relative immunoproteasome activity was increased by IFNβ or TBK1-CM, but not in the presence of the PSMB8-specific inhibitor ONX-0914 (Fig. 5G and 5H). Finally, we concluded that the reduction of monomeric TDP-43 by IFNβ depends on the immunoproteasome pathway, as treatment with the PSMB8-specific inhibitor ONX-0914 completely abolished the IFNβ-induced reduction of monomeric TDP-43 (Fig. 5I). We confirmed that the concentration of ONX-0914 used in Fig. 5I had minimal effects on the canonical proteasome pathway (Fig. S9).

**Figure 5.**
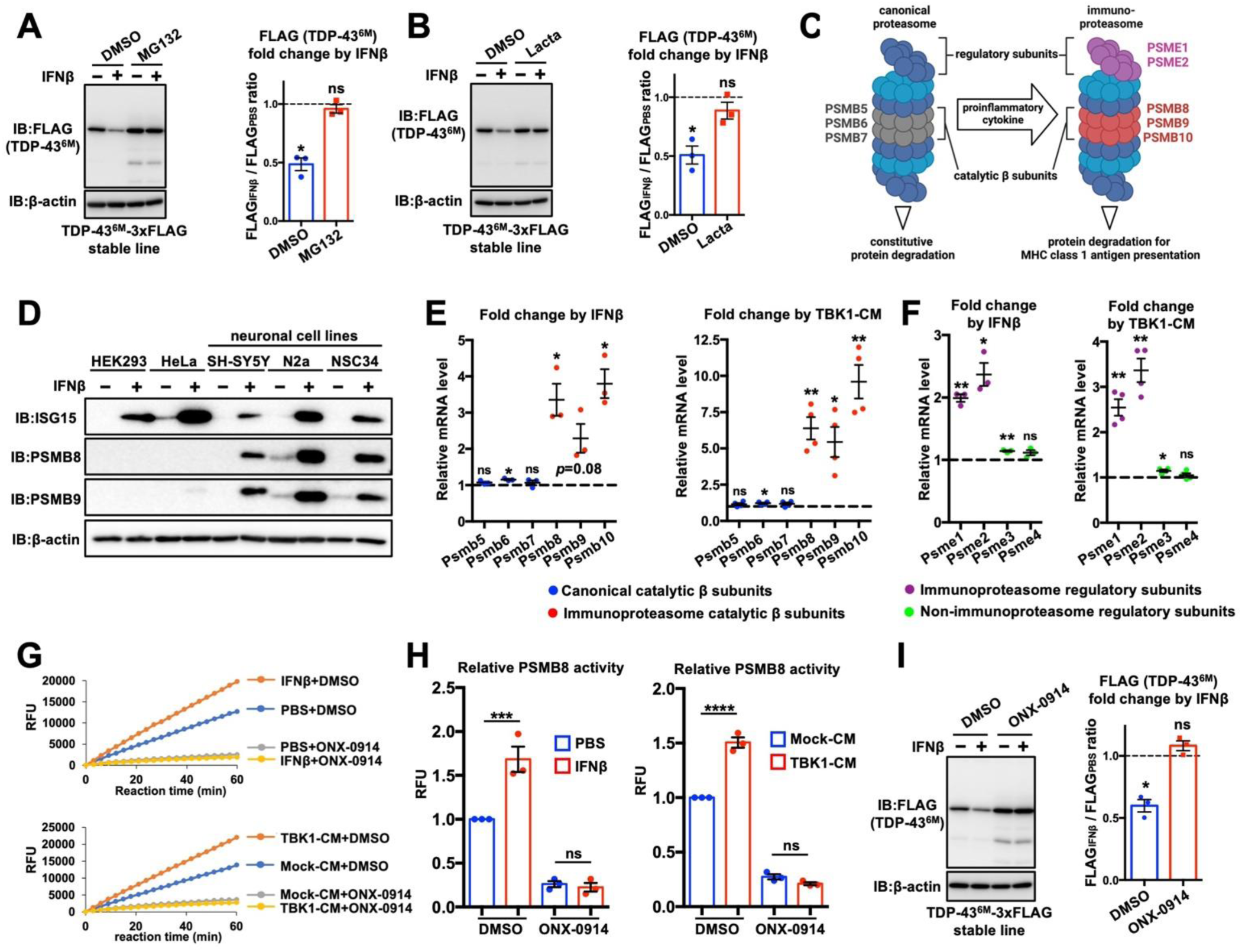
IFNβ upregulates immunoproteasome and promotes degradation of monomeric TDP-43. **A, B.** Proteasome inhibitors abolished the reduction of TDP-43^6M^ by IFNβ treatment. NSC34 cells stably expressing Dox-inducible TDP-43^6M^-3xFLAG were treated with 1.5 μg/ml Dox and either PBS or 10 ng/ml mouse IFNβ for 24h under either DMSO or 10 μM MG132 (A), or 2.5 μM lactacystin (B) treatment. **C.** A schematic illustration of immunoproteasome. **D - F.** IFNβ induced expression of immunoproteasome subunits in neuronal cells. In D, the indicated cell lines were treated with either PBS or 10 ng/ml human/mouse IFNβ for 30h and analyzed by immunoblotting to assess protein levels of immunoproteasome subunits, PSMB8 and PSMB9 (D). ISG15 was assessed to confirm a general response to IFNβ. N2a, Neuro2a. In E and F, Neuro2a cells were treated with either PBS or 10 ng/ml mouse IFNβ, or either Mock- or TBK1-CM for 24h and expression levels of proteasome catalytic β subunits (E) and regulatory subunits (F) were quantified by qPCR. **G, H.** IFNβ induced functional immunoproteasome. Neuro2a cells treated with either PBS or 10 ng/ml mouse IFNβ, or either Mock- or TBK1-CM for 24h were analyzed by immunoproteasome activity assay. Representative kinetic assay results (G) and relative fluorescent unit (RFU) at end points of the assay (H) were shown. ONX-0914-treated samples were prepared as negative controls. **I.** Immunoproteasome-specific inhibition by ONX-0914 abolished the reduction of TDP-43^6M^ by IFNβ. NSC34 cells stably expressing Dox-inducible TDP43^6M^-3xFLAG were treated with Dox and either PBS or 10 ng/m mouse IFNβ for 24h under DMSO or 0.5 µM ONX-0914 treatment. Data are expressed as means ± SEM. ns, not significant. *p < 0.05, **p < 0.01, ***p < 0.001, and ****p < 0.0001.

### Immunoproteasome is upregulated in spinal motor neurons and layer V cortical neurons of wild-type mice with healthy aging

Aging-dependent proteostasis decline is regarded as a major driver of age-related neurodegenerative diseases including ALS and FTD (Hipp *et al*, 2019; Jagaraj *et al*, 2024). Given the potentially protective role of the IFNβ–immunoproteasome pathway in maintaining proteostasis, we next investigated whether type I IFN response and immunoproteasome are upregulated in the spinal cord and brain of mice during healthy aging. In the lumbar spinal cord, representative type I IFN downstream genes, *Ifit1*, *Isg15*, and *Oas1a*, were significantly upregulated in 12-month-old mice compared to younger ages (Fig. 6A). In the cerebral cortex, another type I IFN-responsive gene, *Ifit3*, was significantly increased in 12-month-old mice. However, *Isg15* and *Oas1a* exhibited individual variability and did not show significant changes, suggesting that the elevation of type I IFN response in the cortex at 12 months of age is mild, even if present (Fig. 6B). mRNA expression levels of *Psmb8* and *Psmb9* were also significantly upregulated in both the lumbar spinal cord and cerebral cortex with aging (Fig. 6C and D). Immunostaining revealed that PSMB8 expression was increased primarily in ChAT-positive spinal motor neurons and CTIP2-positive layer V cortical neurons of 12-month-old mice (Fig. 6E and 6F), suggesting an increased requirement for immunoproteasome function in these ALS/FTD-susceptible neurons during aging.

**Figure 6.**
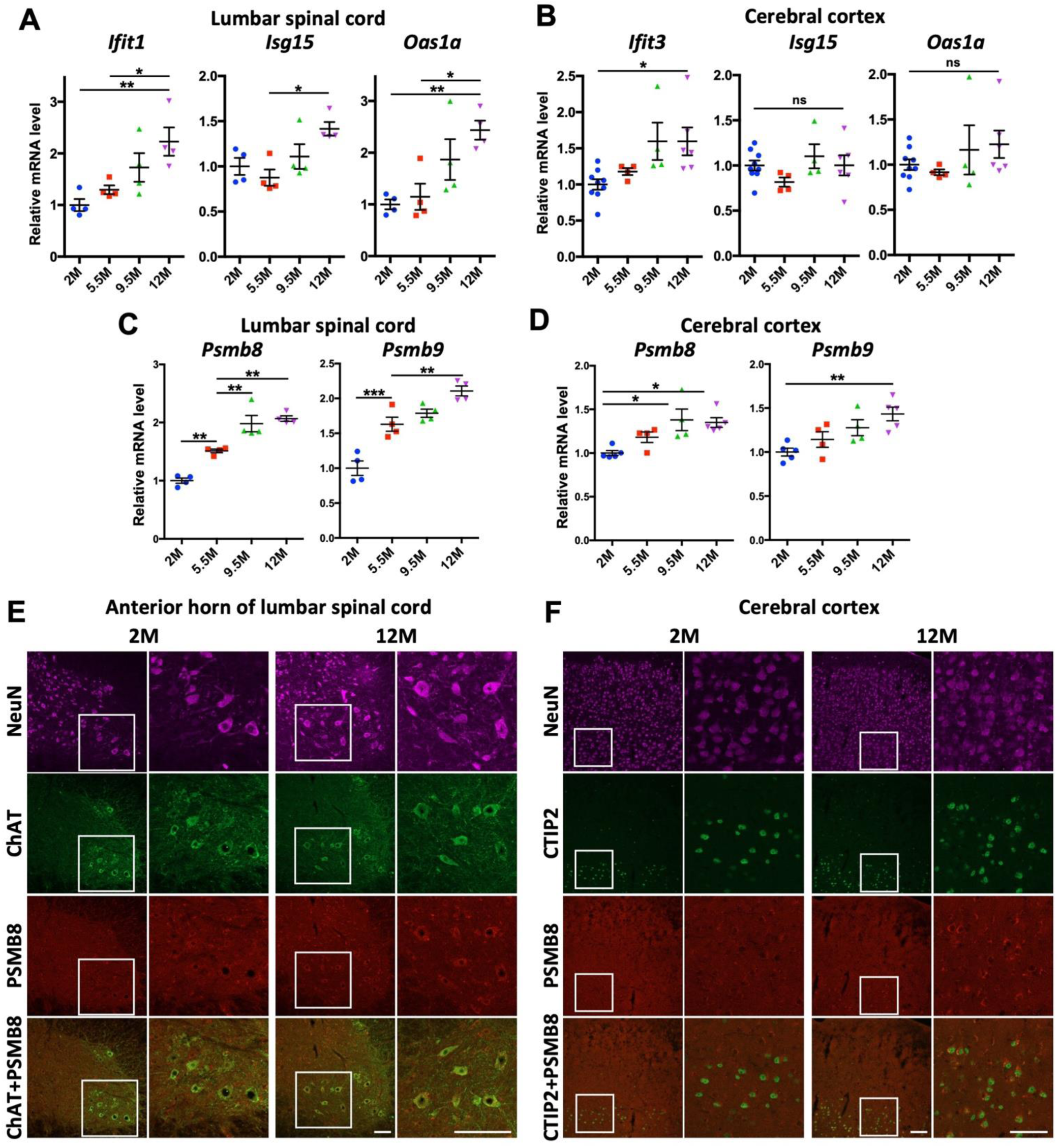
Immunoproteasome increase in ChAT-positive spinal motor neurons and CTIP2-positive layer V cortical neurons with healthy aging. **A, B.** Increased IFN response with aging in lumbar spinal cords and cerebral cortices of wild-type (WT) mice. Total RNAs were extracted from lumbar spinal cords (A) or cerebral cortices (B) of WT mice at the indicated ages. Expression levels of the downstream genes of IFN signaling were analyzed by qPCR. M, month-old. **C, D**. Increased immunoproteasome subunits with aging in lumbar spinal cords and cerebral cortices of mice. Expression levels of immunoproteasome subunits, *Psmb8* and *Psmb9*, were analyzed as with A and B. **G, H.** Increased expression of PSMB8 in ChAT-positive spinal motor neurons and CTIP2-positive layer V cortical neurons in 12-month-old WT mice. NeuN was co-stained as a pan-neuron marker. Scale bar, 100 μm. M, month-old. Data are expressed as means ± SEM. ns, not significant. *p < 0.05, **p < 0.01, and ***p < 0.001.

### Heterozygous deletion of *Tbk1* in an ALS model mouse impairs immunoproteasome induction and exacerbates poly-ubiquitinated protein accumulation at onset stage

Lastly, we investigated the role of TBK1 in immunoproteasome induction in vivo using *SOD1*^G93A^ transgenic ALS model mice (Gurney *et al*, 1994). First, we found that while the canonical proteasome β-subunits, *Psmb6* and *Psmb7*, were downregulated in the lumbar spinal cord of ALS mice at disease onset (3.5 months old) (Fig. 7A), the immunoproteasome β-subunits, *Psmb8* and *Psmb9*, were upregulated (Fig. 7B). These findings, together with the age-dependent upregulation observed in Fig. 6C–F, suggest that immunoproteasome induction may serve as a compensatory mechanism in response to proteasomal demand. Next, we investigated the role of TBK1 in immunoproteasome induction and ubiquitinated protein accumulation in vivo using heterozygous *Tbk1* knockout *SOD1*^G93A^ transgenic mice, which exhibit earlier disease onset compared to normal *SOD1*^G93A^ mice (Brenner *et al*, 2019). Notably, homozygous knockout of *Tbk1* in mice is embryonically lethal (Bonnard *et al*, 2000). We first confirmed that this model exhibited an approximately 50% reduction in *Tbk1* mRNA expression (Fig. 7C). We found that heterozygous deletion of *Tbk1* in *SOD1*^G93A^ transgenic mice led to decreased type I IFN responses, as assessed by reduced expression of *Isg15* and *Oas1a* (Fig. 7D). Immunoproteasome induction in the ALS mice evaluated by *Psmb8* and *Psmb9* was also impaired by heterozygous Tbk1 deletion (Fig. 7E). These results demonstrated that TBK1 plays an important role for induction of type I interferon response and immunoproteasome at the onset stage of ALS. Although we were unexpectedly unable to detect IFNβ in mouse tissues, likely due to its extremely low expression levels (data not shown), it is highly plausible that neurons are the primary source of IFNβ, since phosphorylation of STING at Ser366, a prerequisite for type I IFN induction, was commonly observed in neurons of both aged wild-type and the onset-stage ALS mice (Fig. S10). Finally, to investigate the relationship between impairment of the TBK1–immunoproteasome axis and abnormal protein accumulation, 1% Triton X-100-soluble and -insoluble protein fractions were extracted from the lumbar spinal cords of each genotype and analyzed by immunoblotting. While the levels of an autophagy marker protein p62 did not differ between *SOD1*^G93A^ and *SOD1*^G93A^/*Tbk1*^+/−^ mice, Triton-insoluble polyubiquitinated proteins were more accumulated in *SOD1*^G93A^/*Tbk1*^+/−^ mice, possibly due to impaired induction of immunoproteasome at the onset stage (Fig. 7F). These results suggest that TBK1 deficiency exacerbates abnormal protein aggregation, possibly through impairment of the immunoproteasome pathway in vivo.

**Figure 7.**
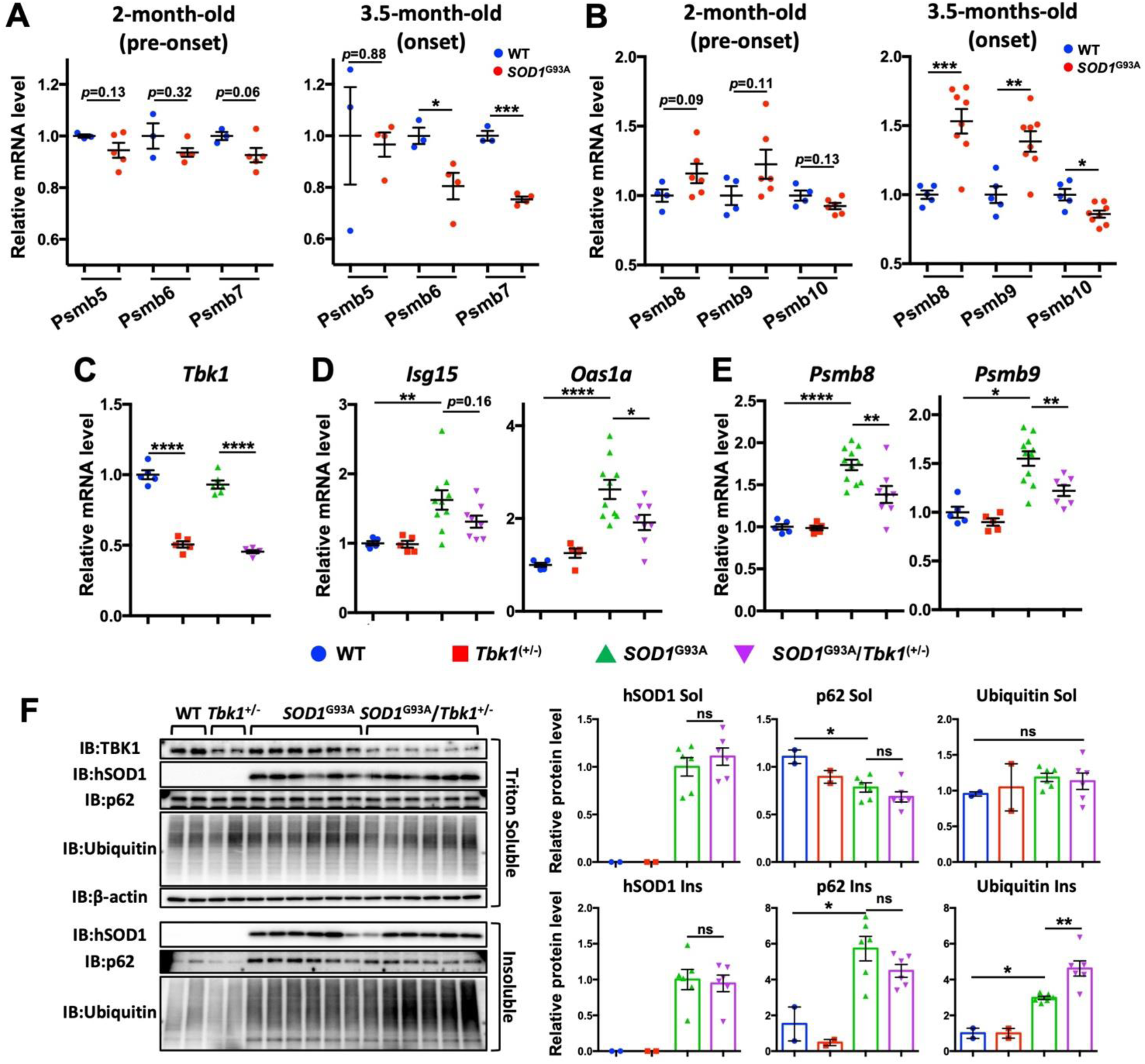
Heterozygous deletion of Tbk1 in SOD1^G93A^ mice causes impaired immunoproteasome induction and exacerbates poly-ubiquitinated protein accumulation. **A.** Canonical proteasome catalytic β subunits, *Psmb6* and *Psmb7*, were downregulated in lumbar spinal cords of *SOD1*^G93A^ mice at onset stage. Total RNAs were extracted from lumbar spinal cords of 2-month-old (pre-onset stage) or 3.5-month-old (onset stage) *SOD1*^G93A^ mice and age-matched wild-type (WT) mice. Relative mRNA expression levels of the indicated genes were analyzed by qPCR. **B.** Immunoproteasome β subunits, *Psmb8* and *Psmb9*, were upregulated in lumbar spinal cords of *SOD1*^G93A^ mice at onset stage. Relative mRNA expression levels of the indicated genes were analyzed as with A. **C-E.** Heterozygous knockout of *Tbk1* in *SOD1*^G93A^ mice impaired the induction of IFN response and immunoproteasome in lumbar spinal cords at onset stage. Total RNAs were extracted from lumbar spinal cords of 3.5-month-old mice with the indicated genotypes and relative mRNA expression levels of the indicated genes were analyzed by qPCR. **F.** Heterozygous knockout of *Tbk1* in *SOD1*^G93A^ mice increased 1% TritonX-100-insoluble poly-ubiquitinated proteins in lumbar spinal cords. Total proteins were extracted from lumbar spinal cords of 3.5-month-old mice with the indicated genotypes, fractionated into 1% TritonX-100-soluble and -insoluble fractions, and analyzed by immunoblotting. Data are expressed as means ± SEM. ns, not significant. *p < 0.05, **p < 0.01, ***p < 0.001, and ****p < 0.0001.

## Discussion

While accumulating evidence has suggested that TBK1 is a key molecule in ALS/FTD, how dysregulation of TBK1 contributes to TDP-43 pathology remains poorly understood. In this study, we demonstrated that TBK1 has the potential to induce immunoproteasome activation via IFNβ signaling, thereby promoting the degradation of aggregation-prone monomeric TDP-43 in vitro (Fig. 8). Importantly, we found that the receptor for IFNβ was downregulated in spinal motor neurons of ALS patients, particularly in TDP-43 pathology-positive motor neurons. We also observed that immunoproteasome expression was induced in ALS/FTD-susceptible neurons of aged wild-type mice and *SOD1*^G93A^ ALS model mice at disease onset. Notably, heterozygous deletion of *Tbk1* in the ALS mice resulted in impaired immunoproteasome induction and increased accumulation of polyubiquitinated proteins. Based on these findings, we propose that the TBK1-IFNβ-immunoproteasome axis represents an inducible and protective mechanism that mitigates proteostatic stress, such as TDP-43 pathology, in neurons. Our results suggest that impairment of this pathway, due to dysregulated TBK1 or reduced IFNAR1 expression, may be one of the mechanisms contributing to the development of TDP-43 pathology in ALS/FTD.

**Figure 8.**
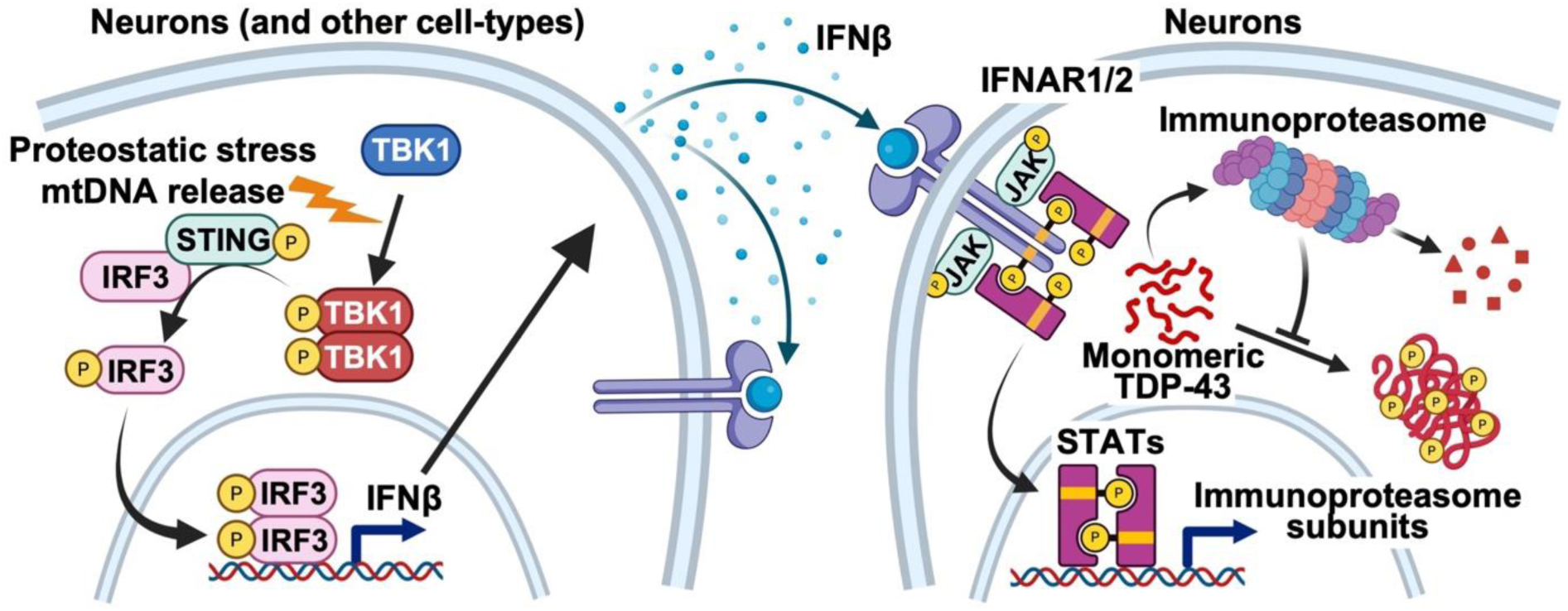
Schematic summary of the TBK1–IFNβ–immunoproteasome axis in TDP-43 clearance. TBK1 is activated by various cellular stresses such as proteostatic stress (Watanabe *et al*, 2023) and mitochondrial DNA (mtDNA) release (Yu *et al*, 2020). Activated phospho-TBK1 phosphorylates STING and IRF3, leading to the induction of IFNβ. IFNβ subsequently upregulates the immunoproteasome in neurons, thereby promoting the degradation of aggregation-prone monomeric TDP-43. In ALS/FTD patients, phospho-TBK1 (Shao *et al*, 2022; Watanabe *et al*, 2023) and IFNAR1 are dysregulated (this paper).

The findings in this study expand the current understanding of TBK1 function beyond its canonical roles in autophagy, endosome, and inflammation, highlighting its potential involvement in proteasome-mediated protein quality control via IFNβ. The TBK1–IFNβ–immunoproteasome pathway possesses two advantages that distinguish it from the autophagy/endosomal–lysosome pathways. First, this pathway depends on proteasome for the degradation of substrates. Misfolded proteins, including misfolded TDP-43, are major substrates of the proteasome, and when not properly eliminated, they can serve as precursors to protein aggregates. Therefore, the proteasome is considered an upstream degradation machinery to maintain proteostasis, and enhancing its activity may represent a more effective strategy for preventing protein aggregate formation, rather than degrading aggregates by lysosome after they have formed. Second, immunoproteasome induction can occur in a non–cell-autonomous manner via IFNβ. This feature enables simultaneous regulation of proteostasis across multiple cells, even when TBK1 is inactive in some of them. In the frontal cortices of C9-FTD patients, phosphorylated TBK1 has been reported to be sequestered into dipeptide repeat protein aggregates, thereby limiting its activity (Shao *et al*, 2022). Under such conditions, neighboring cells with active TBK1 may assist affected cells in mitigating proteostatic stress through IFNβ signaling.

Although TBK1 plays important roles in a wide range of cellular processes associated with ALS/FTD, previous studies have shown that single genetic manipulation of *Tbk1* is insufficient to recapitulate ALS/FTD-related features in mice (Brenner *et al*, 2019; Gerbino *et al*, 2020). Recently, Brenner and colleagues established a TBK1^E696K^ knock-in mouse model, which exhibited progressive motor neuron disease-like symptoms, muscle denervation, and spinal motor neuron loss (Brenner *et al*, 2024). However, TDP-43 pathology was not observed even in the model, indicating the molecular mechanisms underlying TBK1-related ALS/FTD pathogenesis and TDP-43 pathology are complex. This discrepancy between humans and mice may be attributable to the limitation of mice models. Actually, an embryonic stem cell-derived human motor neuron model demonstrated loss of TBK1 activity is enough to cause early-stage motor neuron degenerative phenotypes and mild TDP-43 pathology (Hao *et al*, 2021). In mice models, it seems that TBK1 affects ALS/FTD and TDP-43 pathology as a so-called “second hit”. Sieverding and colleague showed heterozygous deletion of Tbk1 in *TDP-43*^G298S^ transgenic mice worsened neuromuscular junction pathology, although TDP-43 pathology was not altered in the mouse model (Sieverding *et al*, 2021). Another study using a *C9orf72* mouse model of c9FTD-ALS demonstrated that knock-in of disease-linked R228H mutation in *Tbk1* exacerbated phenotypes, neurodegeneration, and pTDP accumulation in the mice (Shao *et al*, 2022). In the present study, we noticed that heterozygous deletion of *Tbk1* in wild-type mice did not alter the expression levels of immunoproteasome subunits and protein accumulation but did in *SOD1*^G93A^ mice (Figure 7). Thus, TBK1 appears to play important roles particularly under disease conditions.

In the present study, we unexpectedly found that lysosomal inhibition did not affect TBK1-mediated clearance of aggregation-prone TDP-43. One possible explanation for this result is that the induced immunoproteasome pathway may act upstream of the lysosomal pathway, and that enhanced proteasome activity alone was sufficient to reduce TDP-43 accumulation, at least under the conducted experimental conditions. It is widely accepted that both the ubiquitin–proteasome system and the autophagy–endosome–lysosome pathway are essential for proteostasis, and the requirement for each degradation system likely depends on various factors such as cell condition and disease stage. Interestingly, it is said that immunoproteasome plays an important role in clearance of defective ribosomal products (DRIPs), which are aberrant, misfolded, or incomplete polypeptides that account for approximately 30% of newly synthesized proteins at physiological conditions (Seifert *et al*, 2010; Schubert *et al*, 2000). The monomeric TDP-43 mutant, TDP-43^6M^, is unable to form its physiological polymeric structure (Afroz *et al*, 2017) and is extremely unstable (Pérez-Berlanga *et al*, 2023), likely due to its misfolded conformation. Thus, it is possible that misfolded proteins associated with neurodegeneration, exemplified by monomeric TDP-43, are preferentially degraded via the immunoproteasome pathway.

While this study suggests a potential protective role for IFN signaling, recent studies have reported that activation of the cGAS–STING pathway is detrimental to ALS progression in both mouse models and iPSC-derived motor neurons (Yu *et al*, 2020; Tan *et al*, 2022). However, the cGAS–STING pathway regulates a wide variety of downstream cytokines, and these studies did not specifically examine the individual roles of each cytokine, including IFNβ. While there is a consensus that proinflammatory cytokines such as TNF, interleukin-1β, and IFNγ exacerbate neurodegeneration through overactivation of glial cells, the role of type I IFNs remains controversial and is likely context-dependent, varying by cell type, disease stage, and disease type. In Parkinson’s disease (PD) model, Ejlerskov and colleagues showed that neuronal IFNβ signaling is protective against PD-like phenotypes and accumulation of PD-related proteins in mice (Ejlerskov *et al*, 2015). The same group also showed that IFNβ is protective against neurodegeneration in PD models by regulating mitochondrial state (Tresse *et al*, 2021). In ALS model, it is said that type I IFN signaling is detrimental because systemic homozygous deletion of *Ifnar1* in *SOD1*^G93A^ mice extended the life span (Wang *et al*, 2011). However, the interpretation of this result is not straightforward because the same study also showed that heterozygous deletion of the gene prolonged the life span more than the homozygous strain. Additionally, the study did not show any results of the onset stage of the strains. As we showed in Figure 7, where *SOD1*^G93A^/Tbk1^+/-^ mice, which show earlier onset than *SOD1*^G93A^ mice (Brenner *et al*, 2019), showed lower type I IFN signaling and more ubiquitinated protein accumulation, there is a possibility that type I IFN signaling at onset stage is protective for ALS. This idea is supported by the study conducted by Nardo and colleagues, where they performed transcriptomic analysis of spinal motor neurons from two *SOD1*^G93A^ strains (C57 and 129Sv) and found that the C57 strain, which exhibits slower disease progression, has higher expression levels of immune response-related genes including type I IFN downstream genes at onset stage (Nardo *et al*, 2013).

This study demonstrated that immunoproteasome is protective against TDP-43 accumulation driven by monomeric TDP-43 in vitro and possibly protective against polyubiquitinated protein accumulation in vivo. However, whether the immunoproteasome is protective in overall ALS/FTD disease course is still open question. A recent study using immunoproteasome subunit knock-out mice demonstrated that immunoproteasome is important for proteostasis in aged mice and lack of immunoproteasome causes epilepsy (Leister *et al*, 2024). Further studies with immunoproteasome deficient ALS/FTD models using animals or iPSC-derived neurons or organoids would be required to address the question.

In conclusion, we found that TBK1 has the potential to suppress proteostatic stress, particularly TDP-43 pathology, through activation of not only autophagy but also the immunoproteasome pathway. Furthermore, we identified IFNβ as an inducer of immunoproteasome in neuronal cells. Although further validation is required, this study is the first to demonstrate that modulating IFNβ signaling and immunoproteasome activity in neurons may represent a potential therapeutic strategy for ALS/FTD.

## Methods

### Antibodies and reagents

The antibodies used in this study were as follows: rabbit monoclonal anti-TBK1 (D1B4) (1:1000; #3504S, Cell Signaling Technology, Danvers, MA, USA), rabbit polyclonal anti-DDDDK-tag (1:4000 for immunoblotting (IB), 1:2500 for immunofluorescence (IF); #PM020, Medical & Biological Laboratories, Tokyo, Japan), mouse monoclonal anti-phosphorylated TDP-43 Ser409/410 (11-9) (1:2000 for IB, 1:2500 for IF; #TIP-PTD-M01, Cosmo Bio, Tokyo, Japan), mouse monoclonal anti-β-actin (1:5000; #A5441, Sigma-Aldrich, St. Louis, MO, USA), rat monoclonal anti-phosphorylated-p62 Ser403 (4F6) (1:500; #D343-3, Medical & Biological Laboratories), rabbit polyclonal anti-LC3 (1:1000; #PM020, Medical & Biological Laboratories), mouse monoclonal anti-Myc-tag (9B11) (1:1000; #2276S, Cell Signaling Technology), rabbit polyclonal anti-RFP (1:1000; #PM005, Medical & Biological Laboratories), mouse monoclonal anti-NeuN (A60) (1:500; #MAB377, Merck Millipore, Burlington, MA, USA), goat polyclonal anti-Choline Acetyltransferase (1:100; #AB144P, Merck Millipore), mouse monoclonal anti-IFNAR1 (H-11) (1:50; #sc-7391, Santa Cruz Biotechnology, Santa Cruz, CA, USA), rat monoclonal anti-CTIP2 (25B6) (1:500; #ab18465, abcam, Cambridge, UK), rabbit polyclonal anti-TDP-43 (1:1000; #10782-2-AP, Proteintech, Rosemont, IL, USA), mouse monoclonal anti-ISG15 (F-9) (1:250; #sc-166755, Santa Cruz Biotechnology), mouse monoclonal anti-PSMB8 (D-2) (1:500 for IB; #sc374089, Santa Cruz Biotechnology), mouse monoclonal anti-PSMB9 (G-2) (1:500; #sc373971, Santa Cruz Biotechnology), rabbit monoclonal anti-PSMB8 (D1K7X) (1:250 for IF; #13635S, Cell Signaling Technology), rabbit polyclonal anti-human SOD1 (raised in our laboratory against a recombinant human SOD1 peptide (aa 24-36)), rabbit polyclonal anti-p62 (1:1000; #PM045, Medical & Biological Laboratories), mouse monoclonal anti-Multi Ubiquitin (FK2) (1:1000; #D058-3, Medical & Biological Laboratories), and rabbit polyclonal anti-phosphorylated STING Ser366 (1:100; #AF7416, Affinity Biosciences, Cincinnati, OH, USA). Alexa Fluor– conjugated and horseradish peroxidase (HRP)-conjugated secondary antibodies were purchased from Thermo Fisher Scientific (Waltham, MA, USA) and Jackson ImmunoResearch Laboratories (West Grove, PA, USA). Reagents were obtained from the following suppliers: Bafilomycin A1 (#B1793, Sigma-Aldrich), Tofacitinib (#S2789, Selleck Chemicals, Houston, TX, USA), recombinant mouse IFNβ (#581302, BioLegend, San Diego, CA, USA), recombinant human IFNβ(#HZ-1298, Proteintech), MG132 (#3175-v, Peptide Institute, Osaka, Japan), Lactacystin (#4368-v, Peptide Institute), Ac-ANW-AMC(#26640, Cayman Chemical, Ann Arbor, MI, USA), ONX-0914 (#16271, Cayman Chemical), BX795 (#SML0694, Sigma-Aldrich), and Doxycycline (#D9891, Sigma-Aldrich).

### Plasmids and siRNAs

Expression plasmids of human TDP-43 (hTDP-43) (WT, 6M, L27/28A, and ΔNLS)-3xFLAG and hTDP-43 (WT and 6M)-mCherry were generated as previously described (Oiwa *et al*, 2023). An expression plasmid of Myc-human TBK1 (WT) was generated as previously described (Watanabe *et al*, 2023). Based on this, ALS/FTLD-linked mutations of TBK1 were introduced by site-directed mutagenesis. For the establishment of stable lines expressing Dox-inducible hTDP-43 (WT or 6M)-3xFLAG, pTRE-Tight-hTDP-43 (WT or 6M)-3xFLAG-SV40-Puro^R^ was generated by subcloning cDNAs of hTDP-43(WT or 6M)-3xFLAG and the SV40 early promotor–puromycin resistance gene– polyadenylation signal sequence from pPur vector (#631601, Takara Bio, Shiga, Japan) into pTRE-Tight vector (#631059, Takara Bio). Predesigned 27mer Dicer-Substrate Short Interfering RNAs (siRNAs) specific for *Sting*, *Irf3*, and *Ifnb1* were obtained from Integrated DNA Technologies (Coralville, IA, USA).

### Cell culture and Transfection

Mouse neuroblastoma Neuro2a (ATCC, Manassas, VA, USA), mouse motor neuron–like hybrid cell line NSC-34 (a kind gift from Dr Neil Cashman, the University of British Columbia), HEK293 (ATCC) and HeLa (ATCC), were maintained in Dulbecco’s modified Eagle’s medium (DMEM) containing 4.5 g/l glucose supplemented with 10% (v/v) fetal bovine serum, 100 U/ml penicillin, and 100 μg/ml streptomycin (Thermo Fisher Scientific). Human neuroblastoma SH-SY5Y (ATCC) was maintained in a 1:1 mixture of DMEM and Ham’s F-12 (DMEM/F12) (Thermo Fisher Scientific) supplemented with 10% (v/v) FBS, penicillin, streptomycin, and ITS-G supplement (Fujifilm Wako Pure Chemical Corporation, Osaka, Japan). For immunofluorescence, Lab-Tek II 4- or 8-well chamber slides (Thermo Fisher Scientific) and Falcon 4- or 8-well culture slides (Corning, Corning, NY, USA) were coated with 40 μg/ml atelocollagen (KOKEN, Tokyo, Japan) at 37 °C for 30 minutes, after which cells were cultured on the coated slides.

In the co-culture experiments, Neuro2a cells transfected with the indicated plasmids were detached using 0.25% (w/v) trypsin–1 mM EDTA solution 6 hours after transfection and replated onto atelocollagen-coated slides or culture plates in the indicated combinations at a 1:1 ratio, using a differentiation medium (DMEM containing 4.5 g/L glucose, supplemented with 2% (v/v) FBS and 2.5 mM N,N-dibutyladenosine 3’,5’-phosphoric acid (dbcAMP) (Nacalai Tesque, Kyoto, Japan).

In the experiments using conditioned media, the culture medium of Neuro2a cells transfected with either mock or TBK1 plasmids was replaced with fresh medium 6 hours after transfection to remove residual plasmids, and conditioned media were collected after an additional 36-hour incubation. The collected media were centrifuged at 1,000 rpm for 5 minutes to remove dead cells, and the supernatant was stored at 4 °C for up to 24 hours before use. The culture medium of recipient Neuro2a cells was replaced with the collected conditioned media 6 hours after transfection with TDP-43 (WT or 6M).

Neuro2a and NSC34 cells stably expressing Dox-inducible hTDP-43 (WT or 6M)-3xFLAG were generated through two sequential rounds of transfection and antibiotic selection. The first transfection was performed using linearized pTet-On Advanced vector (#630930, Takara Bio), followed by positive selection with 1 mg/ml G-418 (Nacalai Tesque) for two weeks. The second transfection was then performed using linearized pTRE-Tight-hTDP-43 (WT or 6M)-3xFLAG-SV40-Puro^R^, followed by positive selection with 2 μg/ml puromycin for an additional two weeks.

Plasmid transfection was performed in Neuro2a, NSC34, SH-SY5Y and HEK293 cells using Lipofectamine 2000 (Thermo Fisher Scientific). For HeLa cells, linear polyethyleneimine (PEI) (Polysciences, Warrington, PA, USA) was used instead. 6 hours post transfection, the media were exchanged to fresh ones. siRNA transfection was performed using Lipofectamine RNAiMax (Thermo Fisher Scientific).

### Animals

Transgenic mice expressing human SOD1^G93A^ mutant (B6.Cg-Tg(SOD1*G93A)1Gur/J) were obtained from the Jackson Laboratory (Bar Harbor, ME, USA). The age of 3.5 months was defined as the onset stage based on our two independent studies using this model (Komine *et al*, 2018; Watanabe *et al*, 2024). *Tbk1* heterozygous knockout mice were established from *Tbk1*, *Ikbke*, and *Tnf* triple knock-out mice (#nbio156) obtained from Laboratory Animal Resource Bank, National Institute of Biomedical Innovation, Health, and Nutrition (Osaka, Japan). C57BL/6J mice obtained from CLEA Japan (Tokyo, Japan) were used as wild-type mice. All mice were housed under specific pathogen-free conditions with 12-hour light/dark cycle at 23 ± 1 °C; 50 ± 5% humidity, with free access to food and water. Mice were handled according to the guidelines established by the Animal Care and Use Committee of Nagoya University.

### Postmortem human tissues

Spinal cord specimens from six patients with ALS and four patients with non-ALS neurological diseases (used as controls) were obtained by autopsy with informed consent. Detailed clinical information for all patients is provided in Table S1. ALS was diagnosed according to the El Escorial clinical criteria established by the World Federation of Neurology, along with the pathological presence of TDP-43 inclusions (Brooks *et al*, 2000). All ALS cases included in this study were confirmed as sporadic through detailed assessment of family history. Ethical approval for the collection and use of human tissues was obtained from the ethics committee of Nagoya University.

### Detergent solubility fractionation and immunoblotting

For detergent solubility fractionation, Neuro2a cells or Mouse lumbar spinal cords were collected in ice-cold TNE buffer (50 mM Tris–HCl (pH7.4), 150 mM NaCl, 1 mM EDTA) containing 1% (v/v) Triton X-100, cOmplete EDTA-free protease inhibitor cocktail (Roche Diagnostics GmbH, Mannheim, Germany), and PhosSTOP (Roche), and lysed by brief sonication on ice. After centrifugation at 15,000*g* for 10 min at 4 °C, the supernatants were collected as soluble fraction. The remaining pellets were washed with the lysis buffer twice and resuspended in TNE buffer containing 2% (w/v) sodium dodecyl sulfate (SDS) by brief sonication. After centrifugation at 15,000*g* for 5 min at room temperature, the supernatants were collected as insoluble fractions. The protein concentration of each sample was determined by using a BCA assay kit (Bio-Rad Laboratories, Richmond, CA, USA). An equal amount of total protein was separated by sodium dodecyl sulfate-polyacrylamide electrophoresis (SDS-PAGE) and transferred to a polyvinylidene difluoride membrane (Immobilon-P; Merck Millipore). The membrane was incubated with a blocking buffer (50 mM Tris-HCl (pH7.4), 150 mM NaCl, 0.05% (v/v) Tween-20, and 2% (w/v) bovine serum albumin (FUJIFILM Wako Pure Chemical Corporation), followed by incubation with primary antibodies diluted in the blocking buffer at 4 °C for at least 6 hours. Bound antibodies were detected with horseradish peroxidase-conjugated secondary antibodies and Immobilon Crescendo Western HRP substrate (Merck Millipore). Images were obtained by LAS-4000 mini (Cytiva, Tokyo, Japan) or LiminoGraph Ⅲ right (ATTO, Tokyo, Japan) and analyzed by Multi-Gauge (Cytiva) or Fiji software (Schindelin *et al*, 2012).

### Immunofluorescence

Cells were plated onto the atelocollagen-coated slides, fixed with 4% (w/v) paraformaldehyde for 20 min at room temperature, and permeabilized with TBS (50 mM Tris-HCl (pH 7.4), 150 mM NaCl) containing 0.5% (v/v) Triton X-100 for 30 min. After blocking with blocking buffer (TBS containing 0.3% (v/v) Triton X-100 and 5% goat or donkey serum) for an hour, cells were incubated with primary antibodies diluted in the blocking buffer overnight at 4 °C. Following three washes with wash buffer (TBS containing 0.3% (v/v) Triton X-100), cells were incubated with Alexa-conjugated secondary antibodies and 1 µg/ml DAPI diluted in the blocking buffer for 2 hours at room temperature. Finally, the samples were mounted with Fluoromount/Plus (Diagnostic BioSytems, Pleasanton, CA, USA).

Mice were deeply anesthetized and transcardially perfused with PBS for 5 minutes, followed by perfusion fixation with 4% (w/v) paraformaldehyde in 0.1 M phosphate buffer for 10 minutes. After an additional overnight immersion fixation in the same fixative, brains and spinal cords were transferred to 30% (w/v) sucrose in PBS and incubated for at least 2 days. Tissues were then embedded in Tissue-Tek OCT compound (Sakura Finetek, Tokyo, Japan) and frozen at –80L°C until use. Cryosections of 20 μm thickness were prepared using a cryostat (Leica Biosystems, Wetzlar, Germany) and immunostained in the same manner as cultured cells. For PSMB8 staining using the rabbit monoclonal D1K7X antibody, antigen retrieval was performed using HistoVT One at 70L°C for 20 min. For IFNAR1 staining, Can Get Signal immunostain Immunoreaction Enhancer Solution A (TOYOBO, Osaka, Japan) was used as the reaction buffer.

For human tissues, 3 μm sections were prepared from paraffin-embedded cervical or lumbar spinal cord samples. Prior to staining, deparaffinization and antigen retrieval were performed using HistoVT One (Nacalai Tesque) at 90L°C for 40 minutes. Immunostaining was performed in the same manner as cultured cells and mouse tissue sections, except that PBS was used instead of TBS containing Triton X-100 in all procedures. Prior to mounting, lipofuscin autofluorescence was quenched using TrueBlack Lipofuscin Autofluorescence Quencher (Biotium, Fremont, CA, USA). Fluorescent intensity of IFNAR1 was quantified using Fiji software.

All images were obtained using a confocal laser scanning microscope (LSM-900; Carl Zeiss AG, Oberkochen, Germany) and the equipped software (ZEN; Carl Zeiss AG).

### RNA isolation and qPCR

Total RNA was extracted using TRIzol Reagent (Thermo Fisher Scientific) according to the manufacturer’s instructions. cDNA was synthesized from 100 - 500 ng of total RNA using the PrimeScript™ RT reagent Kit (Perfect Real Time) (TaKaRa Bio).

qPCR was performed using TB Green Premix Ex Taq^TM^ II (Tli RNaseH Plus) and Thermal Cycler Dice Real Time System II (both from Takara Bio). The primers used in this study were as follow:

### Immunoproteasome activity assay

The immunoproteasome activity assay was performed as previously described (Kim *et al*, 2022) with minor modifications. Briefly, Neuro2a cells were collected in ice-cold proteasome activity lysis buffer (25 mM Tris-HCl (pH 7.4), 5 mM MgCl₂, 10% (v/v) glycerol, 1 mM freshly prepared ATP, 1 mM freshly prepared DTT, EDTA-free protease inhibitor cocktail) and lysed by brief sonication on ice. The lysates were centrifuged at 15,000 × g for 5 min at 4L°C, and the supernatants were collected. Protein concentrations were normalized using the Bradford assay (Bio-Rad Protein Assay Kit). The samples were preincubated with either DMSO or 10 μM ONX-0914 at 4L°C for 30 min. Then, 10 μl of each sample was mixed with 100 μl of lysis buffer in a black 96-well plate, and the fluorogenic substrate Ac-ANW-AMC was added at a final concentration of 12.5 μM. After incubation at room temperature for 15 min, AMC fluorescence was measured every three min using a microplate reader (Infinite 200 PRO, Tecan Schweiz AG, Männedorf, Switzerland) at 360 nm excitation and 460 nm emission wavelengths.

### Statistical analysis

All semiquantitative immunoblotting data, immunofluorescent images, qPCR data, and immunoproteasome activity assay data were analyzed by one-way ANOVA followed by multiple comparison tests with Sidak’s correction for three or more groups and unpaired *t* tests with Welch’s correction for two groups. Outliers were identified using Grubbs’ test at a significance level of α = 0.05 and excluded from both graphical representation and statistical analysis. All statistical analyzes were performed using GraphPad Prism software (GraphPad Software, La Jolla, CA, USA).

## Acknowledgements

This work was supported by Grants-in-Aid for Scientific Research JP22J223468 (to S.S.), JP18H02740, JP19KK0214 and JP22H00467 (to K.Y.) from the Ministry of Education, Culture, Sports, Science and Technology (MEXT), Japan / Japan Society for the Promotion of Science (JSPS); AMED under Grant Numbers JP22ek0109426, JP24wm0425014, and JP24wm0625301 (to K.Y.). S.S. was supported by the Nagoya University CIBoG WISE program from MEXT. The authors are gratefully thankful to the Center for Animal Research and Education (CARE) at Nagoya University for technical support for animal experiments. All cartoons were created with BioRender.com under license to S.S.

## Author contributions

S.S. and K.Y. designed the study. S.S. performed all experiments and analyzed the data with support from K.O., S.W., and O.K., under the supervision of K.Y. Y.I. and M.K. obtained the patients’ autopsy samples and performed neuropathological and clinical diagnoses. Y.I. prepared spinal cord sections from human autopsy tissues. M.H. provided *Tbk1* knockout mice. S.S. and K.Y. interpreted the data and wrote the manuscript. All authors approved the final manuscript.

## Disclosure and competing interest statement

The authors declare that they have no conflict of interest.

## Supplementary figure legends

**Figure S1. TBK1 overexpression does not affect mRNA levels of transfected TDP-43.**

Neuro2a cells were co-transfected with the indicated plasmids and analyzed by immunoblotting and qPCR. Data are expressed as mean ± SEM. ns, not significant; **p < 0.01.

**Figure S2. Suppression of endogenous TBK1 increases pTDP accumulation.**

**A, B.** Neuro2a cells were co-transfected with the indicated plasmids and either control siRNA or siRNA against *Tbk1*, then analyzed by immunoblotting (A) or immunocytochemistry (B). C, D. Neuro2a cells were transfected with the indicated plasmids and treated with either DMSO or 1 μM BX795 (TBK1-specific inhibitor), then analyzed by immunoblotting (C) or immunocytochemistry (D). Scale bar, 20 μm. Data are expressed as mean ± SEM. ns, not significant; *p < 0.05, **p < 0.01, ***p < 0.001.

**Figure S3. Humoral factors induced by TBK1 ameliorates TDP-43 pathology in cells, related to** Figure 1.

**A.** Neuro2a cells were prepared as illustrated in (Fig. 1D) and analyzed by immunoblotting. **B.** Neuro2a cells were prepared as illustrated in (Fig. 1F) and analyzed by immunocytochemistry. Only images of the TDP-43^6M^ case are shown. The ratio of pTDP-positive to FLAG-positive cells is quantified in the graph. Scale bar, 50 μm. Data are presented as means ± SEM. ns, not significant; **p < 0.01, ***p < 0.001, ****p < 0.0001.

**Figure S4. Suppression of endogenous TBK1 did not affect Ifnb1 expression.**

**A.** Neuro2a cells were transfected with either control siRNA or siRNA against *Tbk1*.

**B.** Neuro2a cells were treated with either DMSO or 1 μM BX795 (TBK1-specific inhibitor). Ifnb1 mRNA expression levels were analyzed by qPCR. Data are presented as means ± SEM. ns, not significant.

**Figure S5. Recombinant IFNβ treatment reduces monomeric TDP-43 in a NSC34 stable line.**

**A.** Expression of TDP-43^6M^-3xFLAG in NSC34 cells stably expressing Doxycycline (Dox)-inducible TDP-43^6M^-3xFLAG was confirmed by immunoblotting. **B.** NSC34 cells stably expressing Dox-inducible TDP-43^6M^-3xFLAG were treated with Dox and either PBS or 10 ng/ml mouse IFNβ for 24 hours or 48 hours, then analyzed by immunoblotting. Data are expressed as mean ± SEM. ns, not significant; *p < 0.05, **p < 0.01.

**Figure S6. Recombinant IFNβ reduces mutant SOD1.**

Neuro2a cells transiently transfected with expression vectors of the indicated human SOD1 vectors were treated with either PBS or 10 ng/ml mouse IFNβ for 42h, fractionated into 1% TritonX-100-soluble and -insoluble fractions, and analyzed by immunoblotting. Data are expressed as mean ± SEM. ns, not significant; *p < 0.05, ***p < 0.001.

**Figure S7. Ubiquitous mRNA expression of IFNAR1 in human.**

Relative mRNA expression levels of IFNAR1 in various human cells obtained from the Human Protein Atlas (http://www.proteinatlas.org/celltype) (Karlsson *et al*, 2021) are shown. Magenta arrows indicate neuronal and glial cells.

**Figure S8. Relative mean fluorescent intensity of IFNAR1 in single MN of each patient, related to** Figure 4.

Relative mean fluorescent intensity (MFI) of IFNAR1 in single motor neuron from 4 controls and 6 ALS patients are shown. Data are expressed as mean ± SEM.

**Figure S9. 0.5 μM of ONX-0914 has little effect on canonical proteasome activity.**

NSC34 cells stably expressing Doxycycline (Dox)-inducible TDP-43^6M^-3xFLAG were treated as indicated and analyzed by immunoblotting. Data are expressed as mean ± SEM. ns, not significant; *p < 0.05, ***p < 0.001.

**Figure S10. STING Ser366 phosphorylation is observed primarily in neurons of aged wild-type mice and onset-stage SOD1^G93A^ mice.**

Representative images of phosphorylated STING (Ser366) immunostaining with NeuN and ChAT in lumbar spinal cords of the indicated mice. Scale bar, 100 μm. M, month-old.

## Notes

### Competing Interest Statement

The authors have declared no competing interest.

