## Supplementary figures and images for "TBK1 eliminates aggregation-prone monomeric TDP-43 through the IFNβ-immunoproteasome pathway"

### Supplementary Figures

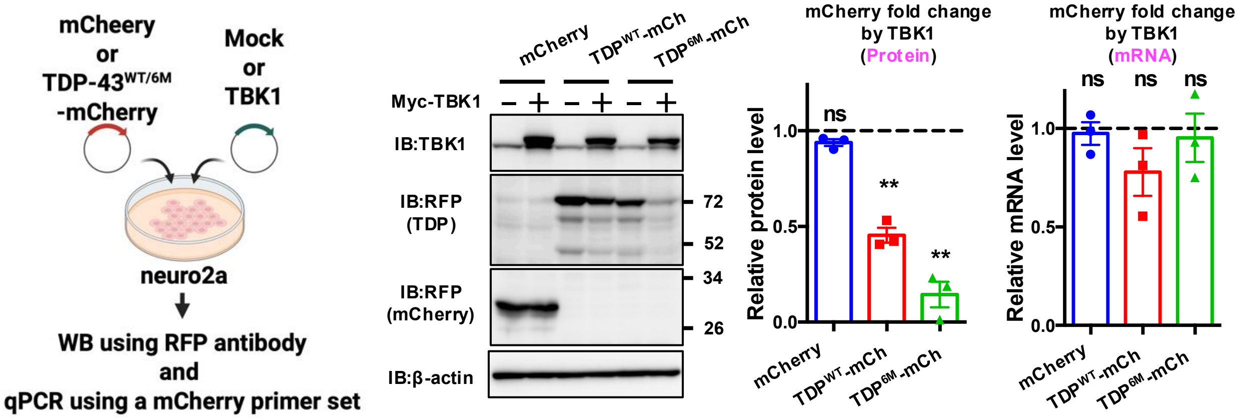

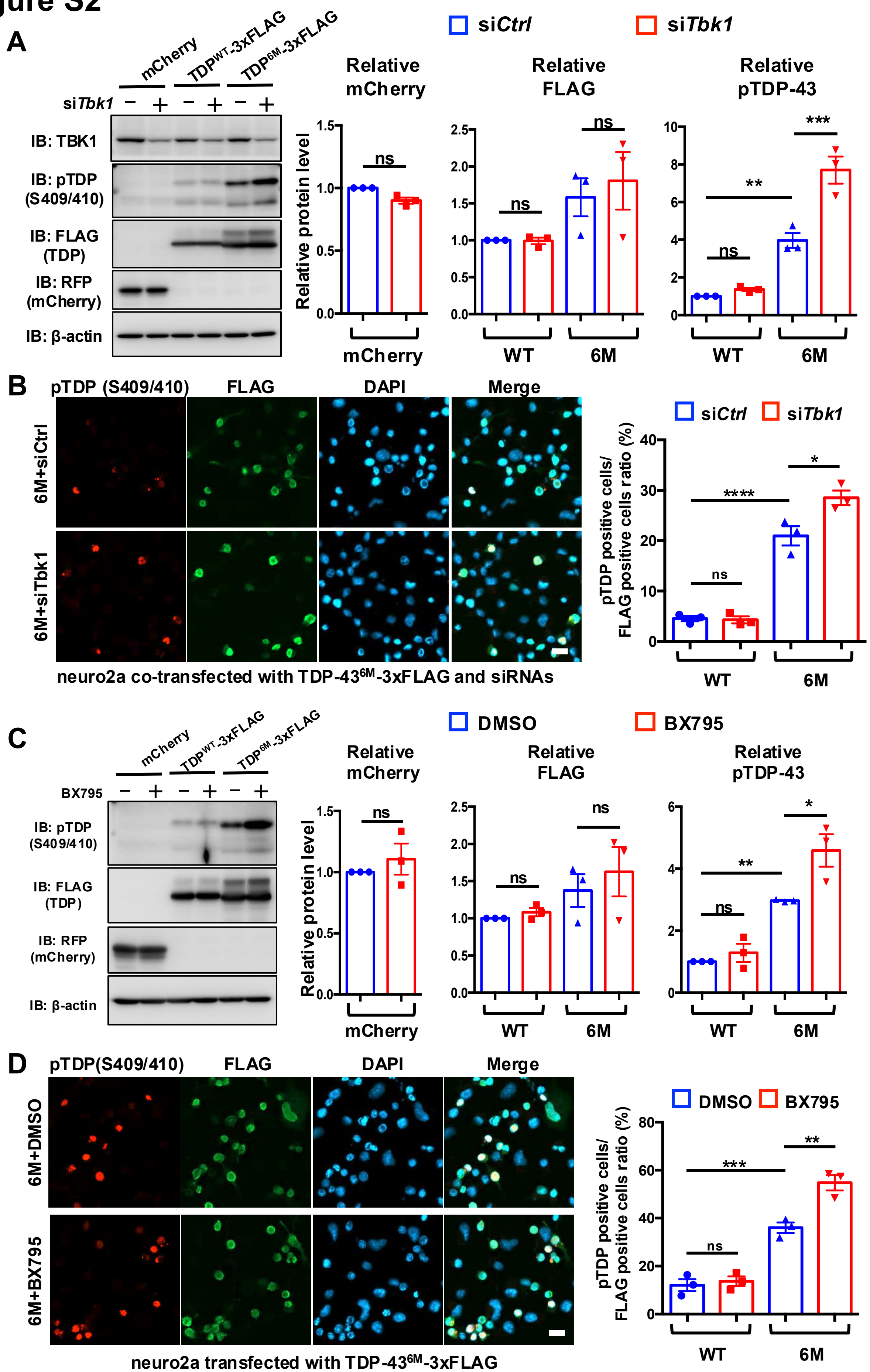

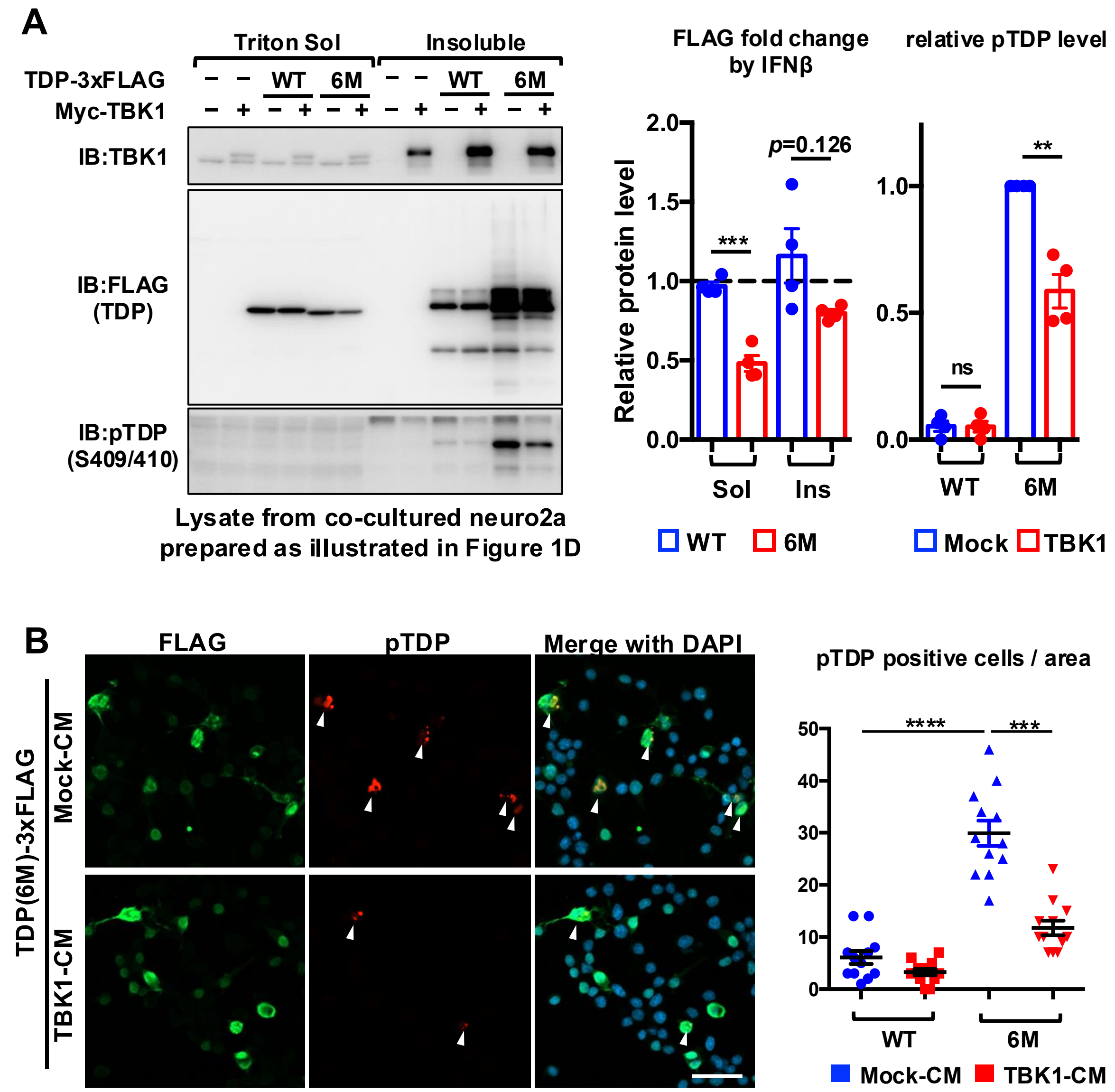

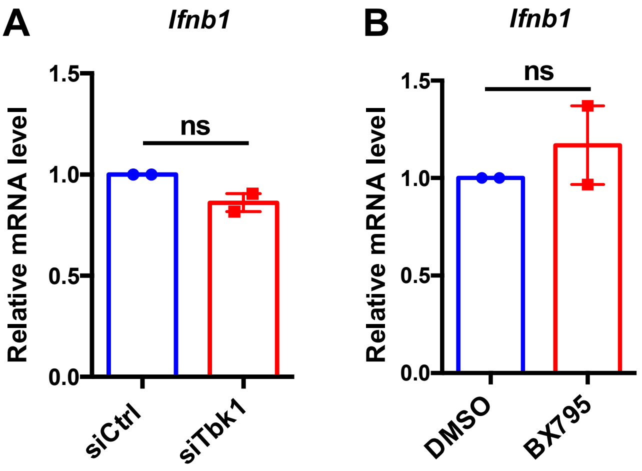

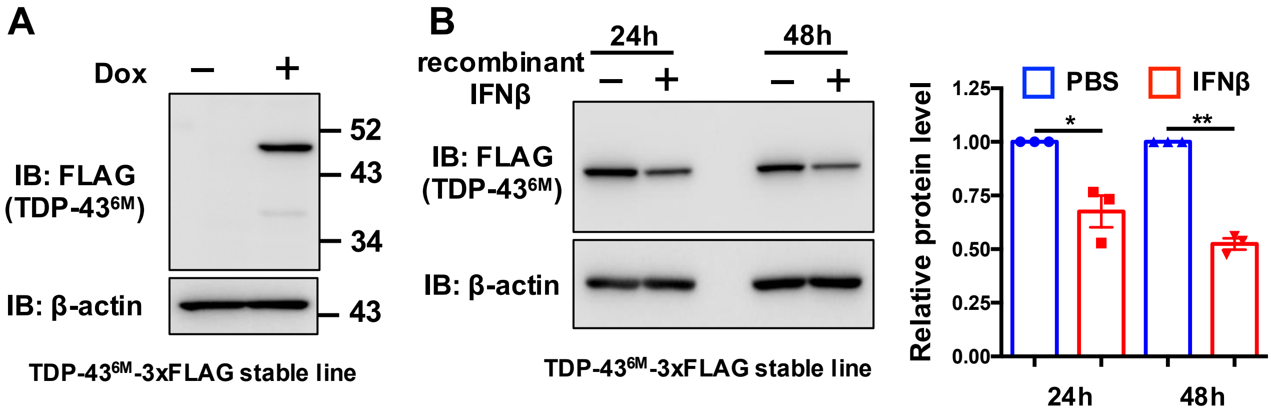

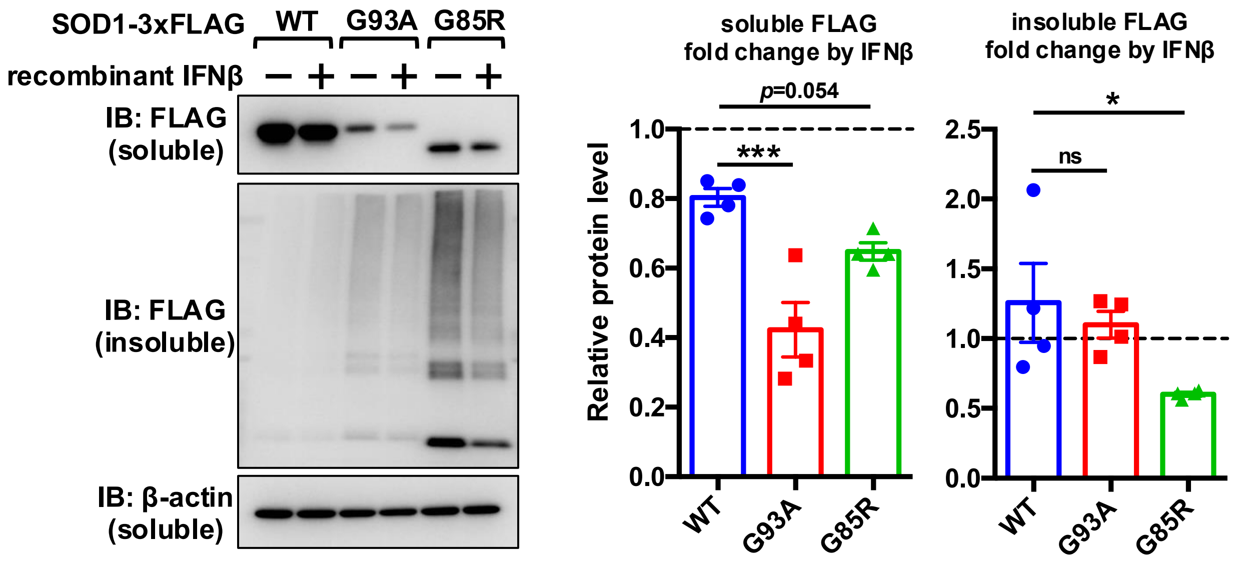

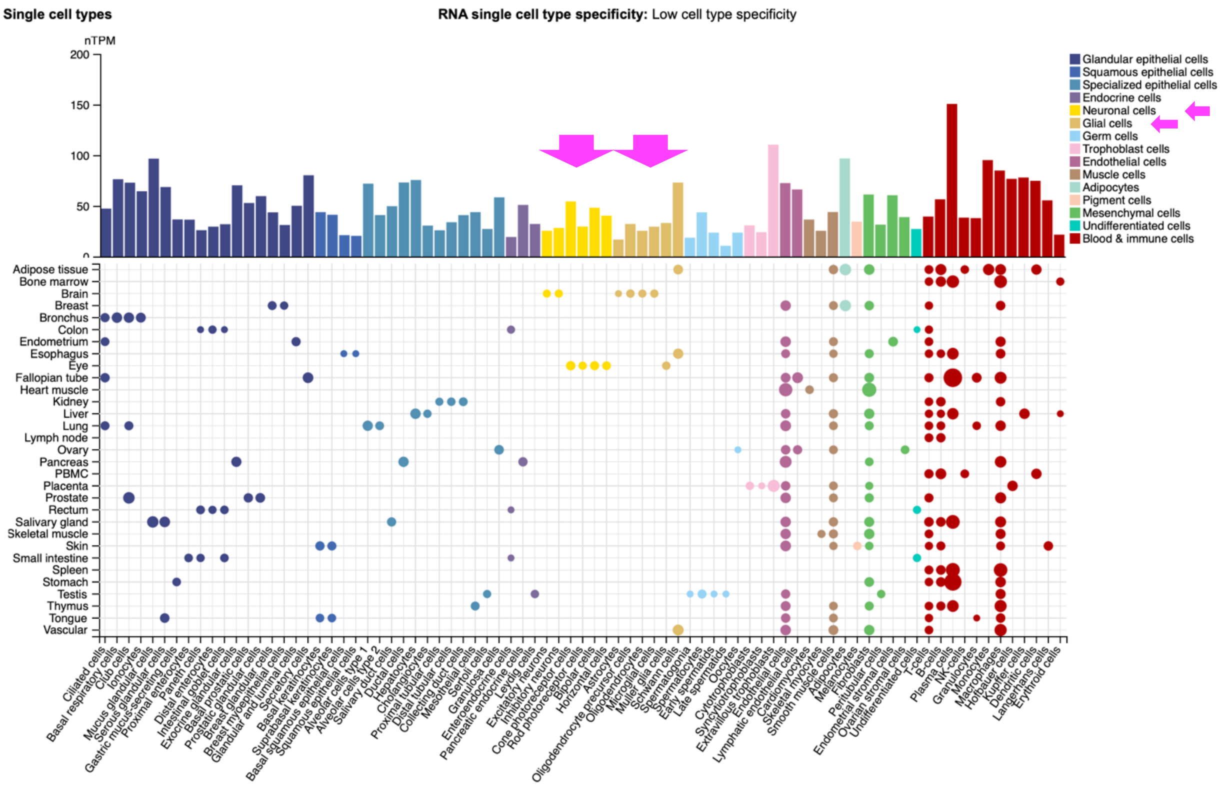

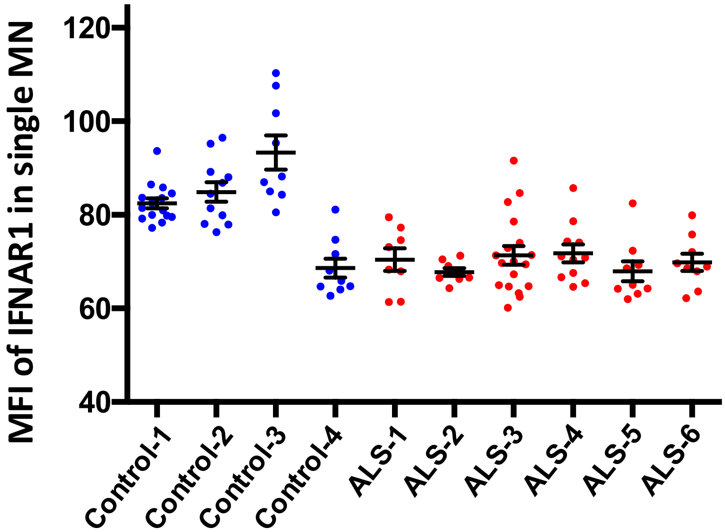

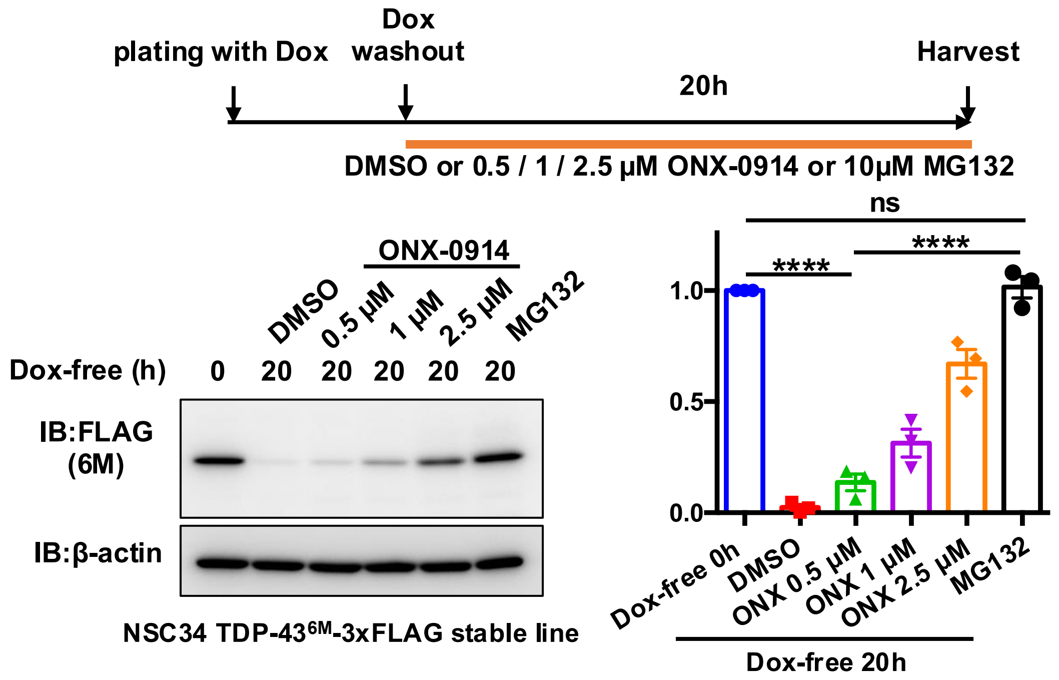

#### Figure S10

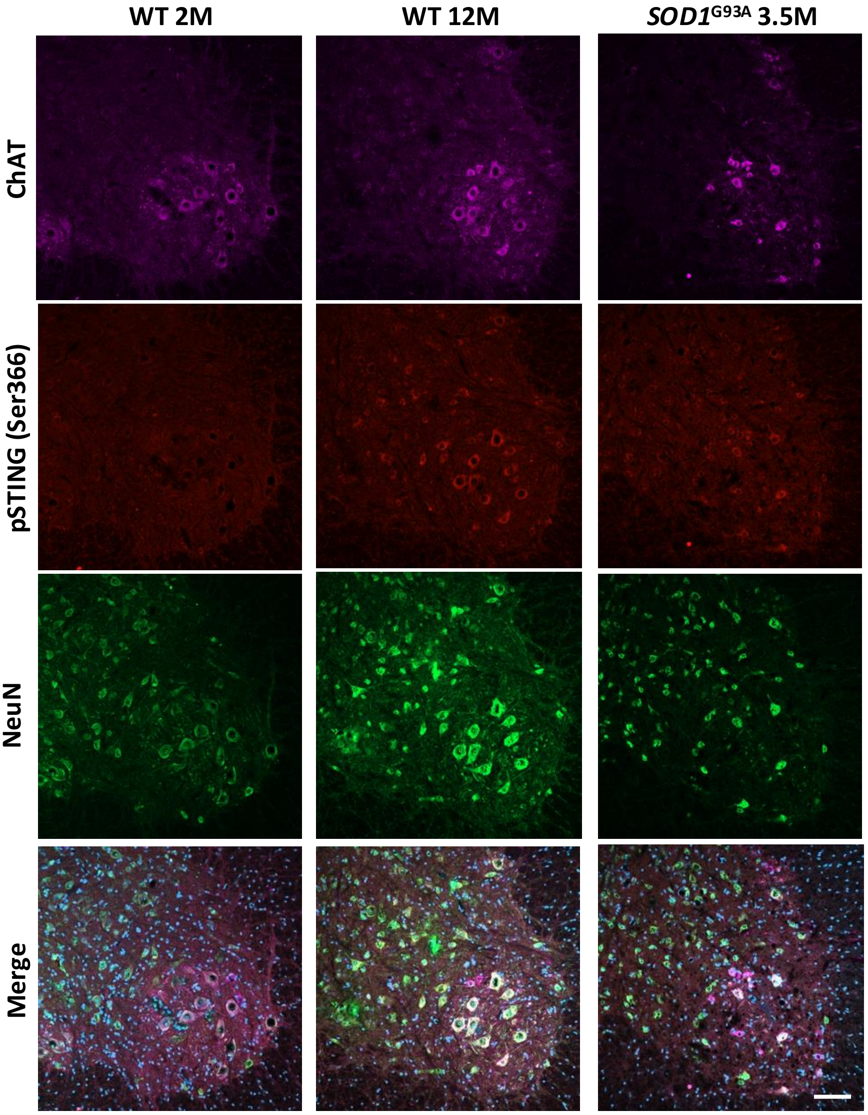
